# Uncovering the transcriptional hallmarks of endothelial cell aging via integrated single-cell analysis

**DOI:** 10.1101/2025.08.18.669055

**Authors:** Sarah Dobner, Lisa Kleissl, Fanni Tóth, Robert Paxton, Irena Yordanova, Kveta Brazdilova, Cedric Vanluyten, Aleksandra Aizenshtadt, Hanna Toth, Jake Burton, Aglaja Kopf, Vera Belyaeva, Katharina Ferencevic, Dunja A. Al-Nuaimi, André F. Rendeiro, Christoph Bock, Joanna Kalucka, Stefan Krauss, Selma Osmanagic-Myers, Laurens J. Ceulemans, Georg Stary, Abdel Rahman Abdel Fattah, Laura P.M.H. de Rooij

## Abstract

Endothelial cells (ECs) are critical regulators of vascular function and exhibit specialized, organ-specific roles across tissues. During aging, these cells become dysfunctional, resulting in increased susceptibility to cardiovascular disease and its associated mortality. While single-cell transcriptomics studies have revealed extensive endothelial heterogeneity across tissues and conditions, a comprehensive atlas of human EC transcriptomes over the course of the adult human lifespan is still lacking. Here, we present the Human Aging Endothelial Cell Atlas (HAECA), a harmonized single-cell transcriptomic compendium of over 375,000 ECs from 12 human tissues throughout adulthood. Using HAECA, we identified age-associated transcriptional shifts, including a decline in angiogenic gene expression in venous ECs and widespread alterations in extracellular matrix (ECM)- and mechanotransduction-associated pathways. We validated these findings in aging human skin and further uncovered a p21-linked transcriptional program in ECs, confirmed in both *in vitro* and *in vivo* models and linked to cellular senescence. Together, our study provides a high-resolution transcriptome reference across spatial as well as temporal axes of the human endothelium.

## Introduction

Endothelial cells (ECs), which line the interior surface of all blood vessels, are central regulators of vascular tone, angiogenesis, permeability, and hemostasis, making them essential for maintaining cardiovascular homeostasis [1]. These cells are highly specialized, adapting to the functional demands of their resident tissues [2]. This is illustrated by their crucial role in maintaining the blood–brain barrier, enabling leukocyte trafficking in lymphoid tissues, or facilitating filtration in the kidney, underscoring the dynamic and organotypic role of the endothelium in human physiology [3–5].

Aging introduces profound alterations to the endothelium, including oxidative and nitrosative stress, impaired angiogenic capacity, and structural remodeling of the vascular wall [6–8]. These changes compromise endothelial repair and regeneration, contributing to a spectrum of chronic conditions such as cardiovascular disease, metabolic syndrome, and neurodegeneration, which all represent leading health burdens in aging populations [9–11]. Importantly, the vasculature is a primary sensor of age-related systemic and circulatory cues, and among the first systems in our body to respond to and be affected by systemic as well as tissue-specific aging-associated factors [12–14]. Moreover, endothelial dysfunction may even precede overt (age-associated) disease [15–18], suggesting that understanding the aging trajectory of ECs may inform early disease detection and prevention strategies.

Advances in single-cell omics have opened new avenues to dissect these aging processes at unprecedented resolution. Especially single-cell transcriptomics studies have uncovered a remarkable degree of heterogeneity among blood vessel ECs, revealing distinct molecular profiles associated to their tissue-specific functions and responses to disease [19–27]. Aging has also been linked to shifts in both cell type composition and gene expression across multiple organs [28–32]. While single studies may be confounded by study-related biases, integration of multiple, independent studies and dataset into one consolidated atlas can bypass these limitations [33]. Indeed, focused integrated atlasing efforts have resulted in detailed insights into the vasculature in several human organs and tissues [25, 27, 34–36]. However, a comprehensive study of single-cell profiles of the blood vessel endothelium throughout the human body and across the human lifespan is still lacking.

Here, we assembled the “Human Aging Endothelial Cell Atlas” (HAECA), a harmonized compendium of over 375,000 human ECs from 12 organs and tissues, covering the adult human lifespan. This comprehensive atlas enables a systematic exploration of endothelial heterogeneity and age-associated transcriptional remodeling across the human endothelium, and a companion interactive web portal (haeca.derooijlab.org) provides easy access to the dataset. To demonstrate the utility of HAECA, we analyzed tissue- and vascular bed-specific aging patterns, revealing a decline in angiogenic signatures in aging veins as well as widespread alterations in mechanotransduction pathways across aging tissue ECs. These findings were validated experimentally in human skin tissue sections. Finally, we used the atlas to identify a p21-associated transcriptional signature linked to EC senescence, which we confirmed both *in vitro* and *in vivo*.

## Results

### An integrated atlas of endothelial cells throughout the human body

To create a detailed atlas of the human vascular endothelium across organs, tissues and ages, we extracted ECs from a total of 34 publicly available sc/snRNA-seq studies (**Figure 1a**; see Methods and **Table S1**). Accompanying metadata was curated and harmonized, to allow for comparisons across tissues, studies, and donor age. Moreover, only cells and nuclei derived from healthy/non-disease human tissue were included, to allow for an analysis of healthy human lifespan.

**Figure 1.**
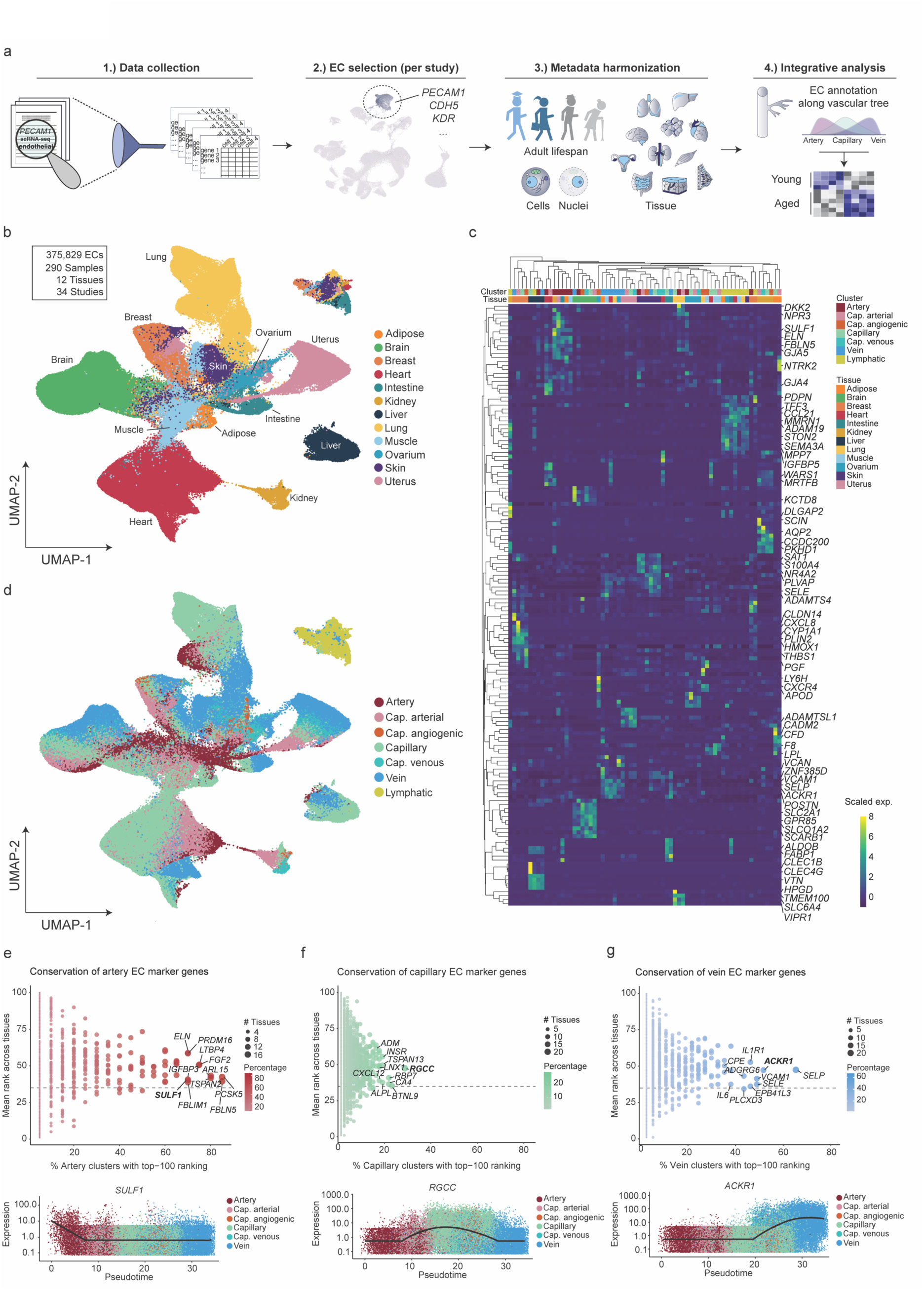
Endothelial transcriptome diversity reflecting organotypic and vascular architecture. **a.** Schematic of study design. **b.** UMAP representation of all ECs incorporated in the final atlas. Color-coding reflects tissue of origin. **c.** Gene-expression heatmap of the top 5 marker genes for each global EC subtype in each tissue. Color scale: yellow, high expression; blue, low expression. **d.** UMAP representation of all ECs incorporated in the final atlas. Color-coding reflects vascular bed subtype. **e-g.** Upper panels: Dot plots of conserved and tissue-specific markers in ECs derived from arterial (e), capillary (f) and venous (g) beds in different tissues. The color intensity and size of each dot represent the fraction (%) and exact number of tissues in which each respective marker gene is found in the top 100 most highly enriched genes, respectively. The top ten most conserved markers are indicated in the plots. Genes in bold are showcased in panels below. Lower panels: expression of representative conserved artery (e), capillary (f), and vein (g) marker genes along the artery-capillary-vein axis in pseudotime. Color-coding reflects vascular bed subtype.

ECs were individually selected from every dataset based on expression of canonical marker genes, including well-described, pan-tissue blood vessel EC (e.g., *PECAM1, CDH5, KDR, FLT1*) and lymphatic EC markers (e.g., *LYVE1, MMRN1, CCL21, PROX1, PDPN*) [19, 27] (**Table S2**). Selected ECs were combined on a tissue-by-tissue basis, followed by data integration, detailed subclustering, and annotation. At this stage, any contaminating (non-EC) and low-quality clusters were removed from the datasets (see Methods).

Our final atlas includes a total of 375,829 ECs, derived from 290 samples and divided across 12 tissues (**Figure 1b, Figure S1a, Table S3)**. Canonical EC markers were robustly detected across tissues, and especially for *CDH5*, *FLT1* and *EGFL7* we observed little tissue-specific heterogeneity (**Figure S1b**). Before integration, a major source of transcriptional variation reflected the sampling method (i.e., sequencing of cells versus nuclei) (**Figure S1c**). We corrected for this technical effect using anchor-based integration, which revealed the underlying tissue-specific transcriptional diversity while preserving biological variation between samples (**Figure 1b**).

We next annotated our ECs according to the major vascular bed subtypes commonly found along the vascular tree (i.e., arteries, capillaries or microvascular ECs, veins, and lymphatic ECs) based on expression of canonical marker genes (**Figure S1d-e, Table S4**). Subclusters with a mixed artery-capillary or vein-capillary transcriptome were annotated as capillary-arterial and capillary-venous, respectively. These clusters likely represent arterioles and venules, respectively, as described previously [19]. Capillary ECs expressing known tip cell markers (e.g., *CXCR4, APLN, ESM1*) [37, 38] were annotated as capillary-angiogenic. These major vascular bed subtypes are annotated in our atlas as “global EC subclusters” and are generally identified in all or most tissues (**Figure S1d-e**). In agreement with previous EC atlasing data in mouse [19], we consistently identified gene expression signatures that could distinguish between ECs derived from different tissues as well as vascular bed subtypes (**Figure 1c, Table S4**). Moreover, again in line with previous reports in human and mouse [19, 27], we could observe transcriptional diversity between lymphatic ECs and blood vascular ECs (i.e., all annotated arterial, microvascular, and venous ECs) in the dataset **Figure 1c-d)**.

For blood vascular ECs, we could furthermore observe gradual phenotypic changes (i.e., zonation) along the arterio-venous axis in most tissues, moving from arteries and arterioles to capillaries, then to venules and veins (**Figure 1d**). To assess the extent to which this zonation is conserved across tissues, we next removed major tissue-specific transcriptional variation in our global atlas (see Methods). We observed that the remaining axis of EC heterogeneity aligned with our global EC subclusters, and arterio-venous zonation became even more apparent (**Figure S1f**). This supports the idea that vascular tree hierarchy is, at least in part, organized by a conserved transcriptional program across tissues.

### Diversity of blood vascular and lymphatic ECs across tissues

Given the known complexity of tissue architecture and the associated specialization of the vascular endothelium, we further subclustered our global EC subtypes. The resulting “fine EC subclusters” enable detailed interrogations of EC subtype diversity across human tissues (see haeca.derooijlab.org and **Table S4**).

Using these detailed EC subtype annotations, we next defined a gene-level representation of the vascular hierarchy by identifying markers that robustly characterize specific positions along the vascular tree (e.g., arterial, venous, or capillary) across tissues. Such conserved markers enable precise identification of distinct EC subtypes (e.g., arterial ECs) throughout the human body, independent of tissue context. To identify these markers, we calculated the top 100 most highly enriched genes for each fine EC subcluster in our atlas. For every EC subtype corresponding to a specific part of the vascular tree, we then computed the mean rank of each gene across clusters and tissues (e.g., all arterial subclusters in all tissues; **Table S5**). This ranking approach allowed us to infer, for each gene, its placement within the vascular hierarchy, and to identify robust cross-tissue markers for each global EC subtype.

Supporting our annotations, this analysis revealed that the previously validated large artery marker *SULF1* [27] was highly conserved in arterial ECs **(Figure 1e, Figure S2a**). Trajectory analysis furthermore revealed the enrichment of this marker at the far arterial end of the vascular tree (**Figure 1e, Figure S2b**). Additional genes frequently detected in arterial ECs across tissues included *FBLN5, PCSK5, ARL15, TSPAN2, FGF2, LTBP4, ELN, FBLIM1, IGFBP3,* and *PRDM16* (**Figure 1e, Figure S2a**).

In line with reports in murine ECs [19], in most tissues, capillary EC clusters were more abundant than arterial, venous, or lymphatic subclusters (**Figure S1e**). As a result, the expression of human capillary markers exhibited greater variability across tissues (**Figure 1f**). Among the most consistently enriched markers were *RGCC*, *CA4*, *BTNL9*, and *RBP7* (**Figure 1f, Figure S2c**). Taking *RGCC* as a representative marker, we confirmed its expected enriched between the arterial and venous ends, i.e., the capillary or microvascular part, of the inferred vascular tree trajectory (**Figure 1f**). Although the expression pattern of the identified conserved capillary genes was heterogeneous across microvascular subclusters, at least one subcluster with strong and selective expression of these markers could be identified in nearly every tissue examined (**Figure S2c, Table S5**). Besides these conserved markers, and in line with literature, we also revealed highly tissue-specific microvascular clusters, including the well-described liver-associated liver sinusoidal ECs (LSECs) [39], lung-associated aerocytes and general capillaries [34, 40], and kidney-associated glomerular ECs [41], each associated with a highly distinct gene expression signature (**Table S4,S5**).

Venous EC markers could also be robustly detected across organs and tissues, with *SELP, ACKR1, SELE, VCAM1, EPB41L3, IL1R1, ADGRG6, CPE, IL6* and *PLCXD3* among the most venous-enriched markers (**Figure 1g, Figure S2d**). Trajectory analysis furthermore confirmed expression of *ACKR1* (used as a representative marker) at the venous end of the vascular tree (**Figure 1g**). When repeating the analysis with the angiogenic capillaries only, we observed a strong conservation of *TP53I11, PXDN, TNFAIP8L1, CTHRC1, F2RL3, LXN, PGF,* and *TNFRSF4* across tissues (**Figure S2e-f**), all previously reported to be enriched in *APLN*-positive tip-like ECs [25]. Selected conserved angiogenic markers (*F2RL3, PGF*) were furthermore enriched along the capillary-venous end of the inferred vascular tree trajectory (**Figure S2g**), in line with the reported emergence of tip cells from venous ECs [25, 42–45]. Lastly, in agreement with findings in the mouse vasculature [19], lymphatic EC markers showed a robust pattern of conservation across tissues and clusters, with *PROX1, MMRN1, COLEC12, MPP7, SCN3B, DOCK5*, *CHRDL1* and *PKHD1L1* ranking among the top conserved genes across LEC clusters in our dataset (**Figure S2h-i**).

### Inflammaging and cellular senescence are conserved hallmarks of the aging endothelium

To allow for an age-associated analysis of EC subtypes and gene expression signatures, we divided our dataset into distinct aged brackets. For every tissue, the median age of all donors was calculated, and donors were categorized as young or aged (below and above tissue median, respectively) (**Figure 2a, Figure S3a-b, Table S3**). While the relative abundance of global EC subclusters was not found to be significantly different between young and aged subgroups, we observed a trend of decreased angiogenic capillary EC abundance in aged samples, in line with reported microvascular rarefaction with increasing age [8, 46] (**Figure 2b**).

**Figure 2.**
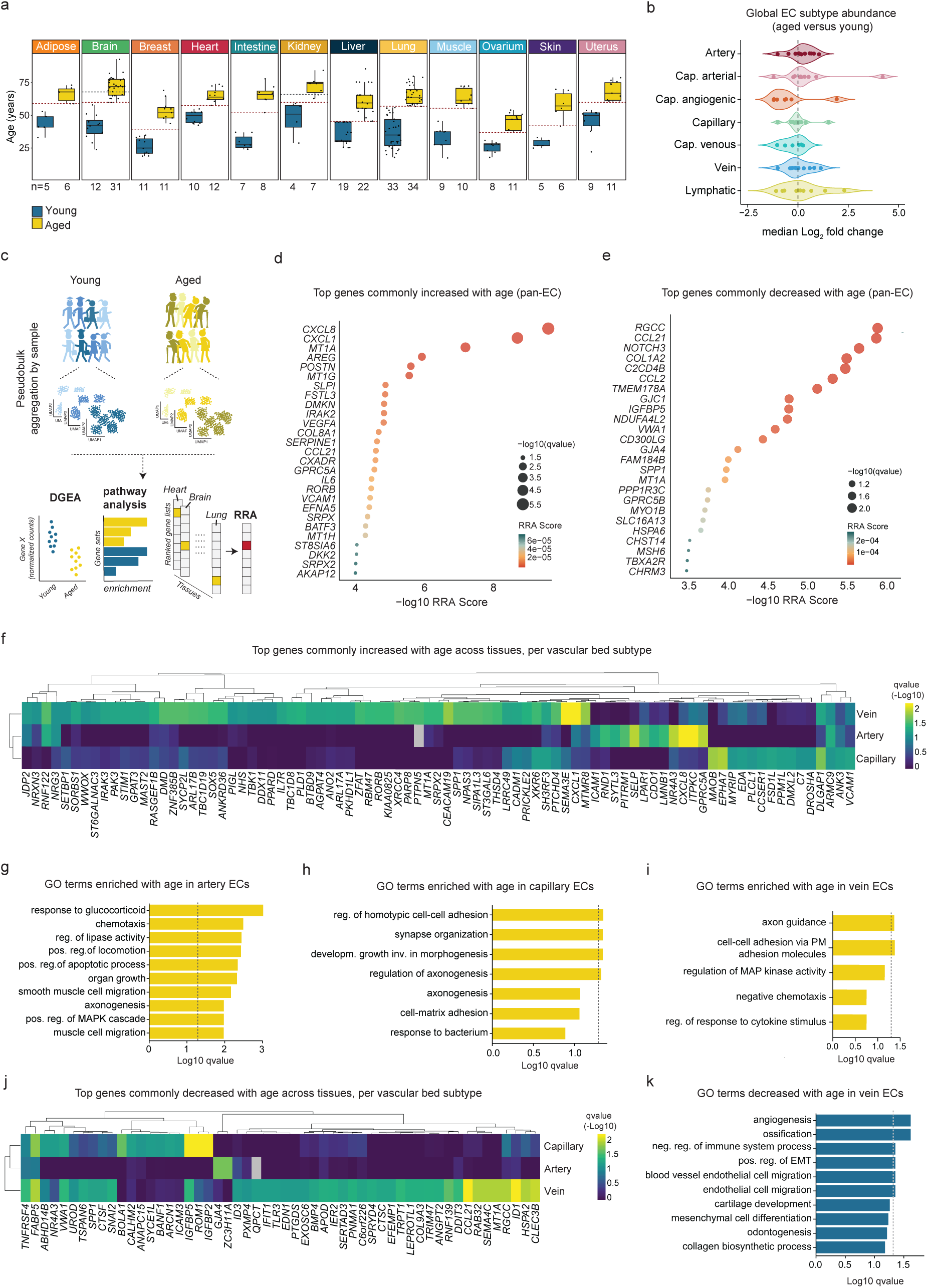
EC aging transcriptome diversity across organs and vascular beds. **a.** Categorization of dataset in age brackets. Young – below tissue median, aged – above tissue median. Red dotted line indicates median age in each tissue. In case of brain and kidney, a cut-off of <60 and ≥60 years old was used to represent young and aged subgroups, respectively, as indicated by grey dotted lines. **b.** Median log2 fold change in abundance (relative fraction per tissue) of global EC subtypes when comparing young and aged subgroups in each tissue. **c.** Schematic of age-associated transcriptome analyses. **d,e.** Dotplot visualization of Robust Rank Aggregation (RRA) results. Top 25 most increased (d) and decreased (e) in aged vs. young ECs across tissues are shown. **f.** Heatmap visualization of markers congruently increased in aged ECs across tissues, per vascular bed subtype. Color scale: yellow, high −log10 qvalue; blue, low −log10 qvalue. **g-i.** GO enrichment analysis, showing top terms enriched in aged artery (g), capillary (h) and vein (i) ECs. Dotted line indicates qvalue ≤ 0.05. **j.** Heatmap visualization of markers congruently decreased in aged ECs across tissues, per vascular bed subtype. Color scale: yellow, high −log10 qvalue; blue, low −log10 qvalue. **k.** GO enrichment analysis, showing top terms decreased in aged vein ECs. Dotted line indicates qvalue ≤ 0.05.

To explore age-associated transcriptome signatures in our dataset, we next aggregated all ECs in a sample-specific manner, to account for variability of biological replicates across studies [47]. The resulting pseudo-bulk dataset was subjected to differential gene expression analysis (DGEA) as well as gene set/pathway analyses within and across tissues, comparing aged and young subgroups (**Figure 2c, Table S6**). Pan aging-enriched or aging-depleted markers (i.e., enriched or decreased in expression in aged ECs across multiple tissues, respectively) aligned with well-described hallmarks of vascular and systemic aging [48–50]. Genes related to immune system regulation, including *CXCL8, CXCL1, IRAK2, CCL21, IL6, VCAM1,* were consistently enriched when comparing aged versus young ECs across tissues (**Figure 2d**). Moreover, the previously reported EC senescence marker *SEMA3A* [51] was among age-increased genes (**Table S6**), suggesting that inflammaging and senescence represent key aging-associated cellular hallmarks of the human aging endothelium independent of tissue context. Among genes robustly decreased in expression when comparing aged to young tissue ECs, we identified angiogenic regulators (*SPP1, NOTCH3*) as well as pan capillary EC markers (*RGCC, CD300LG*) (**Figure 2e**), in line with microvascular rarefaction often observed in the aged vasculature [7, 52]. On the other hand, we identified *VEGFA* as a commonly enriched marker in aged versus young tissue ECs (**Figure 2d**). Whereas a systemic increase of circulating VEGF levels has been shown to counteract age-associated capillary rarefaction in mice [53], evidence suggests that increased VEGFA production may occur during aging in an attempt to compensate for age-associated dysregulated VEGF signaling [49, 54], which would be in line with our results.

Gene set enrichment analysis (GSEA) additionally uncovered molecular signatures that were either consistently enriched or depleted with aging across tissues (pan-EC aging signatures) or changed with aging in a tissue-specific manner (tissue-specific EC aging signatures; **Figure S3c-d, Table S7**). EC aging signatures increased across multiple tissues included those related to known aging-associated hallmarks [48], including cellular senescence and the senescence-associated secretory phenotype (SASP) and inflammation-associated pathways (e.g., TNF-alpha signaling) (**Figure S3c**). Among gene sets significantly decreased in aged versus young ECs across tissues, we found oxidative phosphorylation, EC proliferation, and those associated with MYC targets (**Figure S3d**).

### Extracellular matrix remodeling and decreased angiogenesis in the aging vein EC transcriptome

To explore whether different parts of the vascular tree can be characterized by distinct age-associated transcriptome patterns, we repeated the analyses above with arterial, capillary and venous subsets of the atlas (**Table S8**). Inflammation and migration/chemotaxis-associated changes (e.g., *NR4A3, CXCL8, SELP, ICAM1, ITPKC*) were predominantly observed in aged arterial and capillary ECs (**Figure 2f-i**). In contrast, aged vein ECs were characterized by a distinct set of genes associated with reduced angiogenesis with several angiogenic factors decreased in expression, including *SEMA4C, BMP4, ANGPT2, ID1, ID3*, and *EDN1* (**Figure 2j-k**) [55–60]. Furthermore, among increased genes in aged veins we identified *THSD4* and *SEMA3E*. *THSD4* encodes the secreted Thrombospondin-4 protein, which becomes enriched by TGF-β1 and also plays a role in angiogenesis [61]. SEMA3E has been reported to be either anti-angiogenic or to reduce/normalize angiogenesis in pathological contexts [62–64].

In addition to angiogenic remodeling, aged venous (and capillary) ECs also showed downregulation of *FABP5*, a fatty acid–binding protein critical for endothelial proliferation and chemotaxis [65], as well as decreased expression of *IGFBP2* and *IGFBP5,* encoding insulin-like growth factor–binding proteins. While loss of IGFBP2 may have anti-angiogenic effects [66], IGFBP5 has been shown to accumulate in senescent cells, and its depletion promotes angiogenesis [67, 68]. Aged venous (and capillary) ECs furthermore displayed increased *PTCHD4* expression, a gene associated with survival of senescent cells [69].

Beyond angiogenesis, aged venous ECs showed an upregulation of genes involved in glycosylation, particularly sialylation, across the majority of tissues (**Figure 2f, Table S8**). Notably, expression of sialyltransferases *ST3GAL6* and *ST6GALNAC3* was increased, as well as cell-cell adhesion-associated genes (e.g., *CEACAM19*) (**Figure 2f,i**, **Table S8**). These results suggest that the aged venous extracellular matrix (ECM) environment may be extensively restructured, potentially affecting immune cell trafficking, angiogenesis, and barrier properties.

### ECM-rewiring and altered mechanosignaling in the aging endothelium

Extracellular matrix remodeling is a hallmark of aging and is tightly linked to altered mechanosensing and endothelial function [70]. To explore this further, we analyzed global gene expression changes related to ECM components, glycosylation, and the basement membrane (**Table S7**). Across vascular subtypes, liver, lung, skin, and adipose ECs exhibited the most prominent age-related shifts in ECM- and glycosylation-associated transcripts (**Figure 3a, Figure S4a**). This included increased expression of GALNT family genes (e.g., *GALNT1–3, 10, 11, 15, 17, 18*), which mediate O-glycosylation, as well as multiple sialyltransferases (e.g., *ST3GAL1, ST3GAL6, ST6GAL1, ST6GALNAC3/5, ST8SIA4/6*). In parallel, genes involved in ECM assembly and adhesion (e.g., *LAMA2, LAMA3, COL23A1, TNC, SNED1, THSD4, NTN1, SLIT3*) were enriched in aged ECs. In line with our results above, venous EC clusters (in addition to capillary ECs) showed a prominent age-associated increase for these markers, suggesting broad restructuring of the extracellular landscape with likely consequences for vascular integrity, mechanotransduction, and repair.

**Figure 3.**
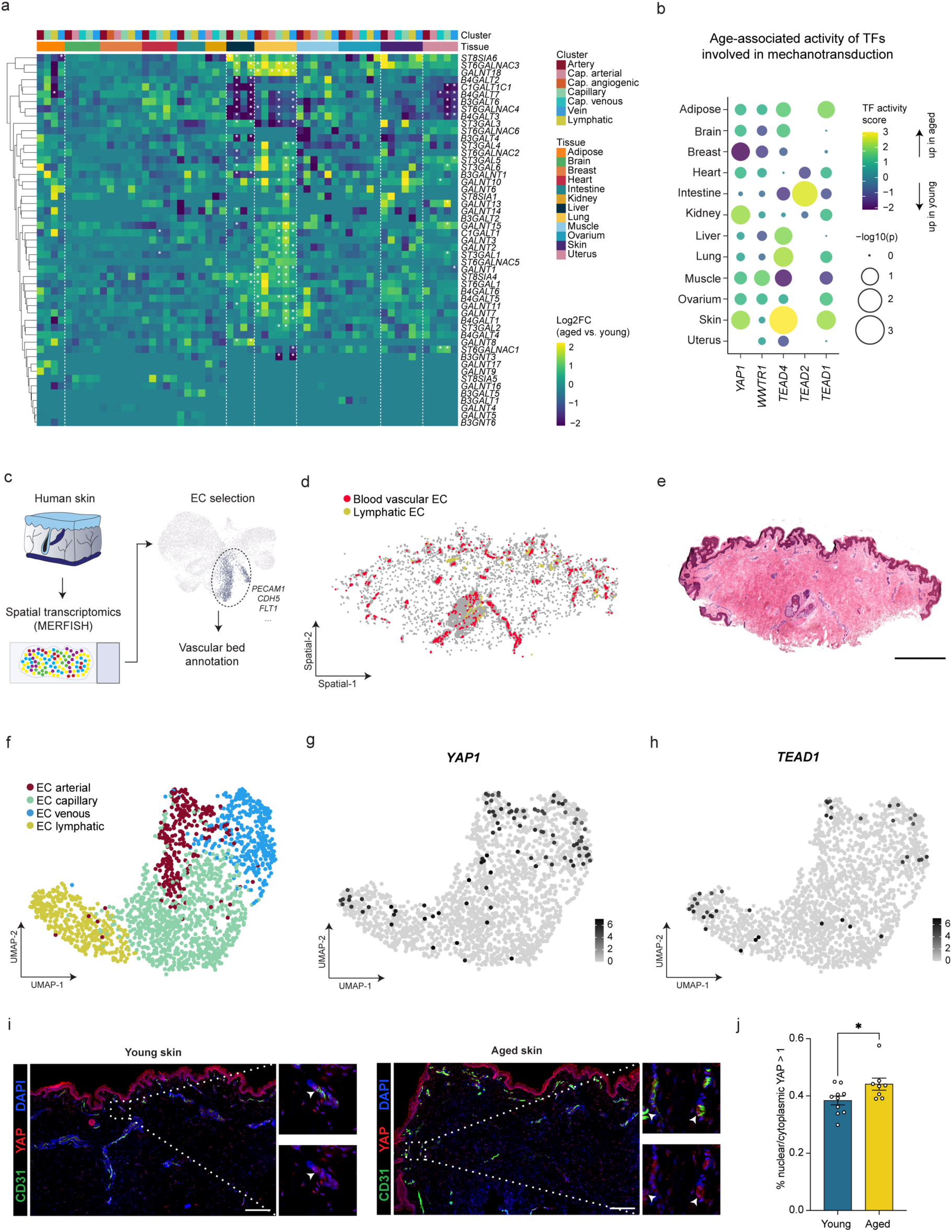
ECM rewiring and altered mechanosensing in the aging endothelium. **a.** Log2 fold change (Log2FC) heatmap of marker genes related to glycosylation, per tissue and vascular bed subtype. Color scale: yellow, increased expression in aged versus young ECs; blue, decreased expression in aged versus young ECs. Asterisks indicate adjusted p-value of 0.1 or lower. **b.** Dot plot heatmap of predicted transcription factor (TF) activities of indicated markers involved in mechanoregulation, across tissue ECs. Color scale: yellow, high activity in aged ECs; blue, high activity in young ECs. **c.** Schematic of spatial transcriptomics analysis. **d.** Representative image of spatially resolved blood vascular (red) and lymphatic (yellow) ECs identified in human skin, as captured via MERFISH spatial transcriptomics. **e.** Hematoxylin and Eosin (H&E) staining of human skin tissue shown in (d). Scale = 500 μm. **f.** UMAP representation of human skin ECs, as captured via MERFISH spatial transcriptomics (n=2). Color-coding reflects vascular bed subtype. **g-h.** *YAP1* (g) and *TEAD1* (h) expression in human skin ECs, as captured via MERFISH spatial transcriptomics. Color scale: back, high expression; grey, low expression. **i.** Representative immunofluorescent images of human skin sections from young and aged subjects, immunostained for YAP (red), CD31 (green), and DAPI (blue). Smaller images are magnifications of the respective boxed areas. Arrowheads indicate nuclei with high YAP signal (within CD31^+^ areas). Scale = 200 μm. **j.** Quantification of the percentage of cells with a YAP nuclear/cytoplasmic ratio > 1 (i.e., indicative of increased mechanical signaling) in young and aged subgroups. Mean ±SEM, unpaired t-test, two-tailed, *p < 0.05, n = 10 and 8 for young and aged groups, respectively.

In the liver, these changes moreover mirrored well-described morphological alterations in the aged sinusoidal endothelium, known as pseudo-capillarization [71]. Aged liver ECs showed upregulation of basement membrane genes *COL4A1* and *COL4A2*, alongside downregulation of the anti-angiogenic factor *COL18A1* (**Figure S4b-d**). At the same time, classical LSEC markers (e.g., *CLEC4G, CLEC4M, STAB1*) were downregulated, consistent with dedifferentiation (**Figure S4d**). However, canonical capillarization markers (*CD34, PLVAP, VWA1*) remained unchanged, suggesting that this molecular profile reflects normal aging rather than fibrosis or inflammation. Aged liver ECs also displayed increased expression of mechanosensitive genes such as *YAP1, WWTR1 (TAZ), PIEZO2*, and *VAV3*, as well as several glycocalyx remodeling genes (*EXT1, CHSY1, ADAM17*), indicating a shift in mechanotransduction and flow sensing (**Figure S4d**). Importantly, these alterations extended beyond microvascular ECs/LSECs, implying a system-wide vascular aging program in the liver.

To further examine whether mechanosignaling pathways are broadly activated across aged ECs in different tissues, we inferred transcription factor (TF) activity from gene expression profiles (decoupleR). Hippo signaling converges on YAP and WW-domain-containing transcription regulator 1 (TAZ), which are transcriptional coactivators that upon activation translocate to the nucleus, where they predominantly interact with the TEAD family of DNA-binding TFs [72]. In line with this, TF activity networks in aged versus young ECs across tissues confirmed increased activity of one or more of the key mechanoregulators YAP1, WWTR1, TEAD1, TEAD2, and TEAD4 across aged tissue ECs (**Figure 3b, Table S9**). While lung and liver ECs showed similar trends towards more active mechanosignaling in aged conditions, this increased activity profile was especially pronounced in the skin endothelium, where YAP1, TEAD4, and TEAD1 were predicted to be significantly more active when comparing aged to young ECs (**Figure 3b, Table S9**).

To confirm endothelial expression of mechanoregulatory markers in a spatial context, we performed spatial transcriptomics on human skin tissue using the MERFISH platform (**Figure 3c**). We identified 1,878 ECs (13.8% of all 13,623 detected cells) and confirmed the expression of global vascular bed subtype markers from our HAECA in this spatial dataset (**Figure 3d–f, S4e-f**). In line with our single-cell atlas, *HEY1, SEMA3G, IGFBP3, CXCL12*, and *FBLN5* marked ECs of arterial nature, *NR2F2* and *ACKR1* identified ECs of venous origin, and *MAF*, *PROX1*, and *CCL21* characterized lymphatic ECs (**Figure S4e-f**), supporting the robustness of our atlas annotations. We moreover detected a relatively large cluster of blood vascular ECs that shared expression of genes including *CLEC14A*, *CD93*, *SPARCL1*, and *HLA-DRA* with arterial and venous ECs, in line with expression patterns of capillary skin ECs in our integrated single-cell dataset (**Figure S4e-f**). We next investigated the expression of genes involved in mechanosignaling and confirmed their increased transcript abundance predominantly in larger vessel ECs (i.e., arterial, venous, and lymphatic EC subtypes) of human skin in both datasets (**Figure 3g-h, Figure S4g**).

To assess the activity of mechanoregulatory factors in ECs in an aging context, we furthermore performed immunofluorescence staining of YAP and CD31 in human skin from young and aged donors (**Table S10**). Here, we observed a subtle yet significantly increased nuclear localization of YAP in the endothelial compartment of tissues derived from aged donors (**Figure 3i-j**), indicating its increased functional activation. These findings are in line with our atlas data, and confirm that mechanotransduction is altered in the aging vascular endothelium.

### A transcriptome signature of endothelial senescence

The accumulation of senescent cells is a well-described hallmark of aging. However, cellular senescence is a highly heterogeneous phenotype, with different cell types and tissues presenting distinct expression patterns of senescence-associated genes and proteins [73]. To deepen our understanding of senescence heterogeneity across cell types, various gene signatures have been curated that can be exploited for a harmonized senescence analysis in (single-cell) RNA-seq data [74–80]. We thus performed a focused GSEA on senescence-associated gene sets in our atlas, and observed that different senescence-associated gene sets showed a variable enrichment pattern across tissues (**Figure S5a, Table S7**). Out of all tissues, aged ECs in the heart were most strongly enriched for senescence gene sets, followed by aged ECs in ovarium, adipose, skin and muscle tissues (**Figure S5a**).

Among the more well-established markers expressed by senescent cells are the cell cycle inhibitors p21^Cip1/Waf^ and p16^Ink4a^ [81]. The number of p21^Cip1/Waf^-positive and/or p16^Ink4a^-positive cells has repeatedly been reported to be significantly increased when comparing aged versus young human or mouse tissues [82–85]. In line with this, we confirmed an increased number of overall p21^Cip1/Waf^-positive nuclei in aged versus young human lung tissues, and a similar trend when specifically analyzing p21^Cip1/Waf^-positive nuclei in the CD31^+^ endothelial compartment of the same samples (**Figure S5b, Table S10**). To identify putative senescent ECs in our atlas, we next investigated expression patterns of *CDKN1A* (encoding p21) and *CDKN2A* (encoding p16). Whereas *CDKN1A* was broadly expressed in ECs in every tissue (average of 42.12% positive cells/tissue), *CDKN2A* was almost undetectable (average of 0.48% positive cells/tissue) (**Figure 4a, Figure S5c**). As reported in other transcriptome studies [86, 87], we did not observe a strong age-associated increase in abundance when comparing the fraction of *CDKN1A*^+^ or *CDKN2A*^+^ ECs with their negative counterparts in young and aged subgroups within most tissues (**Figure S5d**).

**Figure 4.**
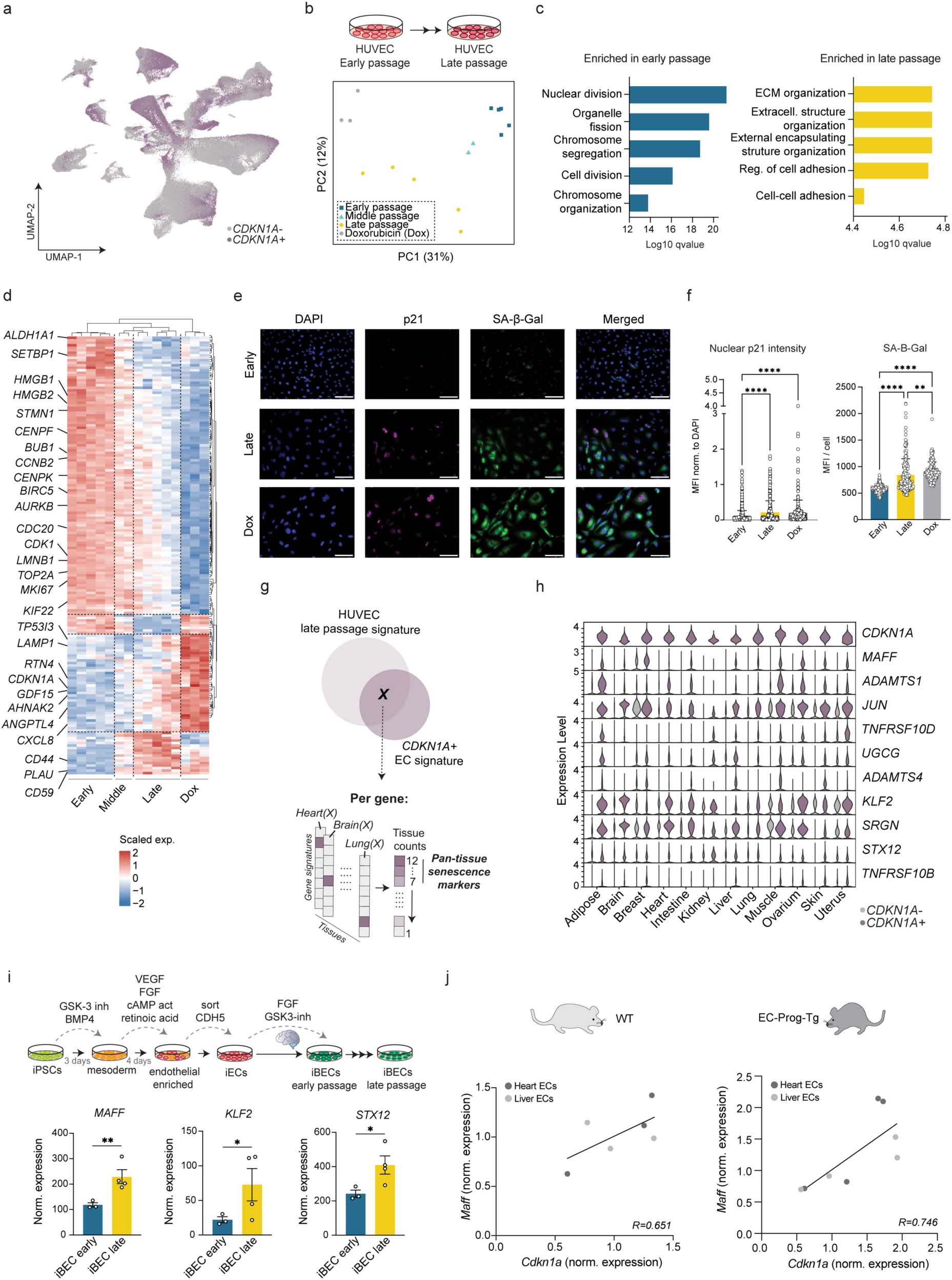
A transcriptome signature of endothelial cell senescence. **a.** UMAP representation of all HAECA ECs. Color-coding reflects *CDKN1A*+ (purple) and *CDKN1A*-(grey) cell fractions. **b.** Principal component analysis (PCA) of early (n = 5), middle (n = 2), late passage (n = 5) HUVECs, as well as early passage HUVECs treated with doxorubicin (n = 3) as analyzed by bulk RNA-sequencing; projection along the two first principal components is shown. **c.** Bar graph visualization of top 5 most enriched GO terms in early (left) and late (right) passage HUVECs. **d.** Heatmap visualization of representative genes enriched at different passage numbers/ conditions of induced cellular senescence. Dashed lines indicate gene clusters associated with the indicated conditions. Color scale: red, high expression; blue, low expression. **e.** Representative images of p21 (magenta), SPiDER-β-gal staining (green) and DAPI (blue) immunofluorescence staining in early and late passage HUVECs, as well as Doxorubicin-induced (Dox) HUVECs. Scale is 100 μm. **f.** Quantification of nuclear p21 staining intensity (left) and SPiDER-β-gal signal intensity (right) in early passage, late passage, and Doxorubicin treated HUVECs (n = 7 each). Mean ±SD, *p < 0.05, by one-way ANOVA. **g.** Schematic of *CDKN1A*-associated signature analysis. **h.** Violin plots of the expression of pan-tissue senescence-associated markers in each tissue. Data is split between *CDKN1A*+ (purple) and *CDKN1A*-(grey) subgroups. **i.** Upper panel: schematic of induced pluripotent stem cells (iPSC) to induced brain-like EC (iBEC) differentiation, and subsequent induction of replicative senescence. Lower panel: differential gene expression analysis between early (n=3) and late passage (n=4) iBECs, for the indicated pan-tissue senescence markers. Wald test, *p < 0.05, **p < 0.01 (adjusted p-values). **j.** Scatterplot of normalized expression of *Cdkn1a* (x-axis) and *Maff* (y-axis) in EC-WT (left) and EC-Prog-Tg (right). Expression was normalized to housekeeping gene *Hprt*. R values are indicated in each plot. n=3 for EC-WT, n=4 for EC-Prog-Tg for each indicated tissue (heart and liver).

To identify additional senescence-associated genes in an endothelial context, we induced human umbilical vein ECs (HUVECs) into replicative senescence via continuous passage. Cells were harvested at different passages and subjected to bulk RNA-sequencing (**Figure 4b**). Doxorubicin-treated HUVECs (passage 3-4), induced into a senescent phenotype after 7 days of treatment, were used as a positive control. When comparing late versus early passage cells, we identified a set of 331 significant senescence-associated genes (p.adj <0.05, Log2FC > 0.35, base mean > 30) (**Table S11**). At early passage, we observed a strong enrichment for genes and Gene Ontology (GO) terms related to cell division, which gradually reduced in expression over the course of continuous passage (**Figure 4c-d)**. Genes related to ECM remodeling and cell adhesion, as well as known senescence markers (e.g., *CD44, GDF15*) became highly expressed in late passage and doxorubicin-induced senescent cultures (**Figure 4c,d**). We verified increased cellular senescence of our late passage and doxorubicin-treated cultures via assessment of senescence-associated beta-galactosidase (SA-β-gal) activity, immunofluorescence stainings with p21, and assessments of nuclear size (**Figure 4e-f, Figure S5e**). Expression levels of *CDKN1A* and *CDKN2A* furthermore gradually increased with passage number in HUVECs, although *CDKN1A* was expressed at considerably higher levels (**Figure S5f**).

Having confirmed that increased *CDKN1A* expression levels are associated with EC senescence *in vitro*, we next compared gene expression profiles of *CDKN1A*+ ECs in our atlas to their respective negative counterparts (within respective tissues). When visualizing the expression of the top 10 most enriched genes in *CDKN1A*+ ECs in every tissue, we observed little tissue-specificity (**Figure S5g**). Indeed, tissue-specific age-associated gene signatures, including decreased expression of LSEC identity markers in liver ECs, were not found to be strongly associated with either the *CDKN1A+* or *CDKN1A-* fraction of ECs (**Table S11**). We confirmed the latter in cultured primary human liver ECs, which did not show decreased expression of canonical LSEC markers *STAB1* and *LYVE1* upon induction of replicative senescence, while *CDKN1A* levels were significantly enriched (**Figure S5h**). These results suggest that *CDKN1A*-associated markers represent a largely conserved transcriptional program across tissue ECs.

Next, we intersected all *CDKN1A*+ EC-enriched genes with the list of 331 HUVEC senescence-associated genes (**Figure 4g**), to further prioritize genes with a putative role in endothelial senescence. Among resulting pan-CDKN1A-associated candidates (i.e., *CDKN1A+* EC-enriched across most tissues, and part of the 331 HUVEC senescence-associated genes) was MAF bZIP transcription factor F (*MAFF*) (**Figure 4h, Table S11**). While MAFF is not traditionally associated with endothelial senescence, it has been implicated in various stress responses and inflammatory pathways [88, 89]. Additionally, MAFF has been identified as a key transcription factor involved in palmitic acid-induced endothelial cell apoptosis, a process relevant to atherosclerosis [90]. Other EC senescence candidates included genes belonging to the ADAMTS family (*ADAMTS1, ADAMTS4*), which may be involved in EC-ECM interactions as well as inflammatory processes. The association of ADAMTS1 and ADAMTS4 with cellular senescence was shown previously in lung tissue and fibroblasts cultures [91], and vascular expression of both was previously reported in the human brain and associated with age-associated neurodegenerative pathologies [92]. Moreover, we identified the AP-1 family member *JUN*, *SRGN* (encoding the proteoglycan Serglycin), the mechanosensing and shear-stress responsive marker *KLF2, TNFRSF10D* and *TNFRSF10B* (encoding TRAIL receptors), and *UCGC* (encoding the Golgi apparatus-residing UDP-Glucose Ceramide Glucosyltransferase) (**Figure 4h**).

To assess whether these genes are strictly associated with p21-associated senescence, or may belong to a broader cellular senescence program, we investigated their enrichment across tissues in EC subpopulations with high (top 25%) and low (bottom 25%) expression scores of two curated gene signatures of cellular senescence (SenMayo [74] and CellAge [93], see Methods and **Figure S6a,c**). When comparing differentially expressed genes between the two subpopulations of ECs, the majority of our senescence candidates were among genes commonly enriched in the SenMayo/CellAge^high^ populations across multiple tissues (**Figure S6b,d, Table S11**), confirming their putative association with cellular senescence in ECs (independent of *CDKN1A* expression). SenMayo predominantly consists of SASP factors (83/125 genes), whereas the CellAge signature is comprised of genes experimentally validated to induce cellular senescence in human cell lines. Whereas both signatures resulted in highly similar expression patterns of our candidate senescence markers in low, middle and high EC populations across tissues (**Figure S6b,d**), their enrichment was considerably higher in the CellAge^high^ setting (in comparison with SenMayo^high^) (**Table S11**), possibly suggesting that their role in senescence may be partially independent of (or go beyond) SASP.

To confirm the association of these putative senescence markers in an alternative, organotypic endothelial context, we differentiated induced pluripotent stem cells (iPSCs) into brain-like endothelial cells (iBECs) (**Figure 4i**). iBECs showed expression of typical blood brain barrier (BBB) endothelial markers at both RNA and protein levels (**Figure S6e-f**). Moreover, key iBEC-associated genes could be mapped to brain ECs in our atlas (**Figure S6g**), confirming their brain-like phenotype. RNA-sequencing furthermore confirmed significantly enriched expression of *MAFF*, *KLF2* and *STX12* in late passage iBECs (**Figure 4i, Table S11**). We moreover observed trends for enriched expression of *Maff*, *Adamts1*, and *Ugcg* in aortic ECs from *Lmna^G609G/G609G^*versus WT mice (scRNA-seq dataset derived from [94]), which recapitulate various features of Hutchinson-Gilford progeria syndrome (HGPS), including increased cellular senescence (**Figure S6h**). When analyzing freshly isolated heart and liver ECs from Prog-Tg (expressing human progerin exclusively in ECs) and WT mice, we furthermore observed that *Cdkn1a* and several of our candidate senescence markers, including *Maff,* were positive correlated (**Figure 4j, Figure S6i, Table S11**). This correlation was strongest in Prog-Tg ECs (**Figure 4j**).

Altogether, by integrating transcriptomic data across tissues, datasets, species, and models systems (*in vitro* and *in vivo*), we uncovered a conserved *CDKN1A*-associated program in ECs with potential relevance for age- and/or pathology-associated vascular senescence.

## Discussion

In this study, we built a comprehensive reference transcriptome resource of EC heterogeneity across the human body and adult lifespan. By integrating over 375,000 high-quality EC profiles from 34 datasets spanning diverse tissues and ages, we captured the complex organization of the vascular endothelium along both tissue-specific and vascular bed-specific axes. Our analysis revealed that, despite strong transcriptomic organotypicity, ECs consistently display conserved transcriptional programs within arterial, venous, lymphatic, and angiogenic compartments across tissues. Especially capillary EC transcriptomes also showed a high degree of tissue-specificity, in line with their dominant role in mediating organ-specific exchange functions and adapting to the unique metabolic and structural demands of each tissue microenvironment [19, 95–97]. Although the expression of conserved capillary marker genes was heterogeneous across capillary subclusters, nearly every tissue contained at least one subcluster with strong and selective expression of these markers. These findings may suggest a recurring transcriptional organization within the microvasculature, where tissue-specific endothelial diversity coexists with shared transcriptional programs associated with capillary specialization throughout the body.

When comparing aged and young subgroups of ECs across tissues, aged ECs consistently showed increased expression of inflammatory mediators and senescence-associated genes, in line with systemic inflammaging and endothelial dysfunction observed during aging. These conserved transcriptomic changes provide strong support for the idea that EC aging involves a shared, tissue-independent program of immune activation and cellular senescence, contributing to loss of endothelial homeostasis.

Among vascular subtypes, venous ECs showed the most pronounced and consistent age-related transcriptomic changes across tissues. These changes include reduced angiogenic signaling, increased expression of pro-senescent and survival-associated genes, and remodeling of the extracellular environment. This is particularly notable given that venous ECs are the origin of angiogenic sprouting [25, 42–45], suggesting that their functional impairment could therefore directly contribute to impaired vascular regeneration and perhaps even capillary rarefaction in aging.

In line with the observed signatures of ECM remodeling in aged ECs, loss of the EC glycocalyx has been associated with advancing age [98]. Recent work has moreover highlighted a structural deterioration of the endothelial glycocalyx during aging in the blood brain barrier [99]. While we also observed age-associated decreases in the expression of several genes encoding galactosyl- or sialyltransferases, a large proportion of them were in fact found upregulated. While speculative, increased sialylation via *ST6GALNAC3* on venous and capillary ECs may reflect a compensatory or age-associated remodeling of the glycocalyx to maintain vascular barrier or signaling function. ST3GAL6 furthermore promotes synthesis of sialyl Lewis X (sLeX), a ligand facilitating leukocyte adhesion, which could contribute to enhanced immune cell-endothelium interactions in aging tissues. In addition, many genes belonging to the GALNT family of proteins, encoding enzymes that regulate the initial step in mucin O-glycan synthesis, were found increased in expression when comparing aged to young ECs. While certain O-glycans were described to become enriched during replicative senescence of human fibroblast cultures [100], O-glycosylation remains rather understudied in the context of aging. However, O-glycosylation-associated genes are often overexpressed in cancer, where they are believed to play a role in altered cell proliferation, migration, invasion, adhesion, as well as immune infiltration [101, 102], suggesting similar functional consequences may occur in an aging context as well. Age-associated altered expression of GALTNs was not homogeneous across tissues in our atlas, with lung, liver, skin and adipose tissue among those with the strongest changes. Our findings may thus open exciting novel avenues for the exploration of tissue-specific EC glycocalyx restoration, potentially with a specific focus on O-glycosylation.

In line with the observed ECM-associated transcriptome changes in aged ECs across tissues, our observations of increased nuclear localization of YAP in aged human skin ECs furthermore suggest elevated endothelial responses to mechanical cues in an aged tissue environment. In the context of the endothelium, activation of YAP/TAZ signaling in ECs was found as a key mechanism of vascular pathology in a HGPS mouse model of accelerated aging [94]. Moreover, increased YAP activity was shown previously in replicative senescent HUVECs [103]. However, age-associated changes in YAP activity are heterogeneous, and endogenous YAP/TAZ activity has been repeatedly reported to decrease with advancing age in non-endothelial contexts (i.e., aged keratinocytes and dermal fibroblasts) [104, 105]. This reduction is however not universal across all cell types, as aging does not seem to significantly impact YAP/TAZ activity in epithelial cells, neurons, or lymphocytes [106]. Increased YAP activity has furthermore been reported in the context of chronic pathologies including lung and liver fibrosis, where Verteporfin (an inhibitor of the YAP-TEAD interaction) was shown to be beneficial in a preclinical context [107]. The same inhibitor was also shown to harbor senolytic capacities [108], further underscoring the potential relevance of YAP/TAZ in an endothelial aging context. Whether our observed YAP activation in aged skin ECs represents a compensatory mechanism to maintain endothelial integrity or a pathogenic trigger promoting senescence, fibrosis, and/or inflammation remains to be elucidated. Future studies will be required to determine the functional consequences of sustained YAP activity in aged ECs across tissues, and to explore whether pharmacological modulation of YAP signaling in the endothelium could offer therapeutic benefit in mitigating age-associated vascular deterioration.

Lastly, our results support the notion that EC aging phenotypes are heterogeneous, comprising a complex mix of adaptive stress responses (e.g., increased VEGFA expression potentially as a compensatory mechanism) to dysfunctional, pro-inflammatory, and anti-angiogenic states. In line with this, we identified *MAFF*, *KLF2*, and *JUN* as upregulated in CDKN1A+ ECs and senescent HUVECs. These genes are also known to be involved in vascular stress responses, mechanosensing, and inflammatory adaptation, not just aging or cellular senescence [88, 90, 109, 110]. In addition, the process of angiogenesis and microvascular sprouting is highly dependent on senescence and the expression/activity of canonical senescence-associated markers (e.g., p21^Cip1/Waf^) [111]. This functional overlap suggests that elevated expression of canonical senescence markers in ECs may not relate explicitly to an age- or pathology-associated terminally arrested state, but may also reflect a context-dependent, transient activation related to vascular remodeling. The same applies to our newly identified CDKN1A-associated transcriptional program, the functional role of which in aging, senescence, and angiogenesis, remains to be elucidated.

## Limitations of the study

We acknowledge that our findings are limited by the respective study, tissue, and age bracket cohort sizes. Further functional validation will increase the robustness of our findings across populations and organs. Sample selection was based on the quality and depth of the metadata reported in each respective study and dataset. Therefore, our results may be biased to unreported age- and tissue-associated confounders (e.g., undocumented comorbidities, lifestyle factors). Age distributions were moreover not similar across all tissues and not always well-balanced, affecting certain comparisons. Some observed EC subclusters may be influenced by sampling methods, such as differences between single-cell and single-nucleus data or differences in tissue digestion protocols. However, our identified subclusters are in strong in agreement with previously published clustering of the vascular endothelial compartment in a range of tissues [19, 27, 34], highlighting the robustness of our integrated dataset. While sex-associated differences in vascular aging are well known [112, 113], such effects were not explored in this study due to the relatively small cohort sizes in each tissue group. Similarly, effects of demographic, exposures, or lifestyle factors (e.g., smoking, ethnicity, medication) could not be explored due to lack of available metadata (in most cases) and the size of our sample cohorts. Stringent gene filtering was applied to ensure robustness of our age-associated analyses across tissues and studies, which may have excluded subtle but biologically relevant signals. Finally, senescence analysis is limited by the heterogeneous nature of senescent cells/ECs and the lack of definitive markers for every tissue and cell type [73, 78, 114]. *CDKN1A*, for instance, may not be universally reliable or specific to senescent ECs and future identification of better, more specific age-associated EC senescence markers will be required to further finetune/complement our findings.

Despite these limitations, and given the current scarcity of pan-tissue atlases focused on healthy human aging, our findings offer important novel insights into the gene expression landscape of the aging vasculature and may inform future efforts regarding the screening and treatment of age-related vascular pathologies.

## Data and code availability

To ensure accessibility, reproducibility and resource value of the HAECA, we made our data available for further exploration via an interactive webtool at haeca.derooijlab.org. Links to the raw data from every HAECA_ID can be found in **Table S1**. Processed .RDS files for every tissue will be made available at Zenodo upon publication (https://zenodo.org/records/16779452). Code used for major age-associated analyses is available at https://github.com/deRooijLab/HAECA. All raw and processed bulk RNA-sequencing data generated during this study will be made available upon publication at GEO.

## Acknowledgments

We thank the Biomedical Sequencing Facility at CeMM for assistance with sequencing, and Stephan Reichl for help and advice. We furthermore thank the Genomics-RNA Core Facility of the Medical University of Vienna for assistance with MERSCOPE sequencing, and Roland Foisner for providing the Prog-Tg mice. Moreover, we thank the Division of Surgery, Inflammatory Diseases and Transplantation, and Department of Gastrointestinal and Paediatric Surgery, of the Oslo University Hospital Rikshospitalet for providing liver tissue samples (Dr. Sheraz Yaqub, Prof. Espen Melum). This work was supported by Angelini Ventures S.p.A. Rome, Italy (S.D., L.K., F.T., R.P., A.F.R., L.d.R.), grants from the Austrian Science Fund (P37301, L.d.R.; FWF P37003, S.O-M; FWF PAT8196024, AR.AF.), a DOC Fellowship of the Austrian Academy of Sciences (OeAW; S.D.), and funding from the Research Council of Norway (262613) and Welcome LEAP Dynamic Resilience program (A.A., K.F., S.K.). L.d.R. is supported by the Clinical Research Group MOTION, Medical University of Vienna, Vienna, Austria – a project funded by the Clinical Research Groups Programme of the Ludwig Boltzmann Gesellschaft (Grant Nr: LBG_KFG_22_32) with funds from the Fonds Zukunft Österreich.

## Supplementary Figures

**Figure S1.**
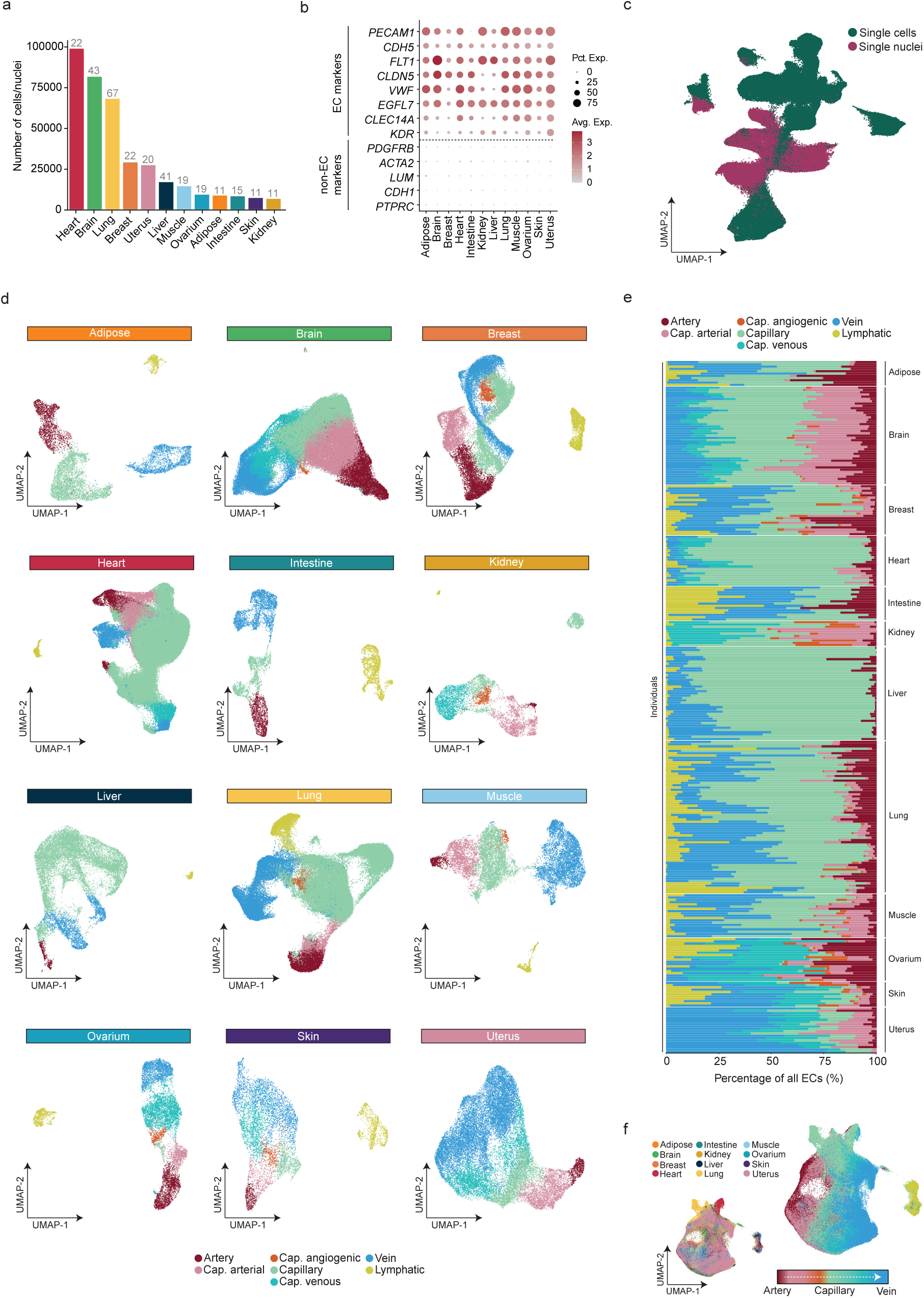
Endothelial transcriptome diversity reflecting organotypic and vascular architecture. **a.** Bar graph visualization of number of endothelial cells/nuclei per tissue, as incorporated in the final atlas. The number of individuals per tissue is indicated above each bar. **b.** Dot plot heatmap of representative EC and non-EC marker genes in every tissue. The color intensity of each dot represents the average level of marker expression, the dot size reflects the percentage of ECs expressing the marker within each tissue. **c.** UMAP representation of all ECs incorporated in the final atlas, without integration by/correction for the sampling method (i.e., sequenced cells/nuclei). Color-coding reflects ECs derived from single cells (green) or single nuclei (maroon). **d.** UMAP representation of all ECs captured for each individual tissue. Color-coding reflects global EC subtypes. **e.** Bar graph visualization of the relative distribution of global EC subclusters in every tissue and individual incorporated in our atlas. Each bar represents a separate individual/sample, x-axis represents the fraction (%) of all ECs in each sample. Color-coding reflects global EC subtypes. **f.** UMAP representation of all ECs, integrated by tissue. Color-coding reflects individual tissues (left) or global EC subtypes (right).

**Figure S2.**
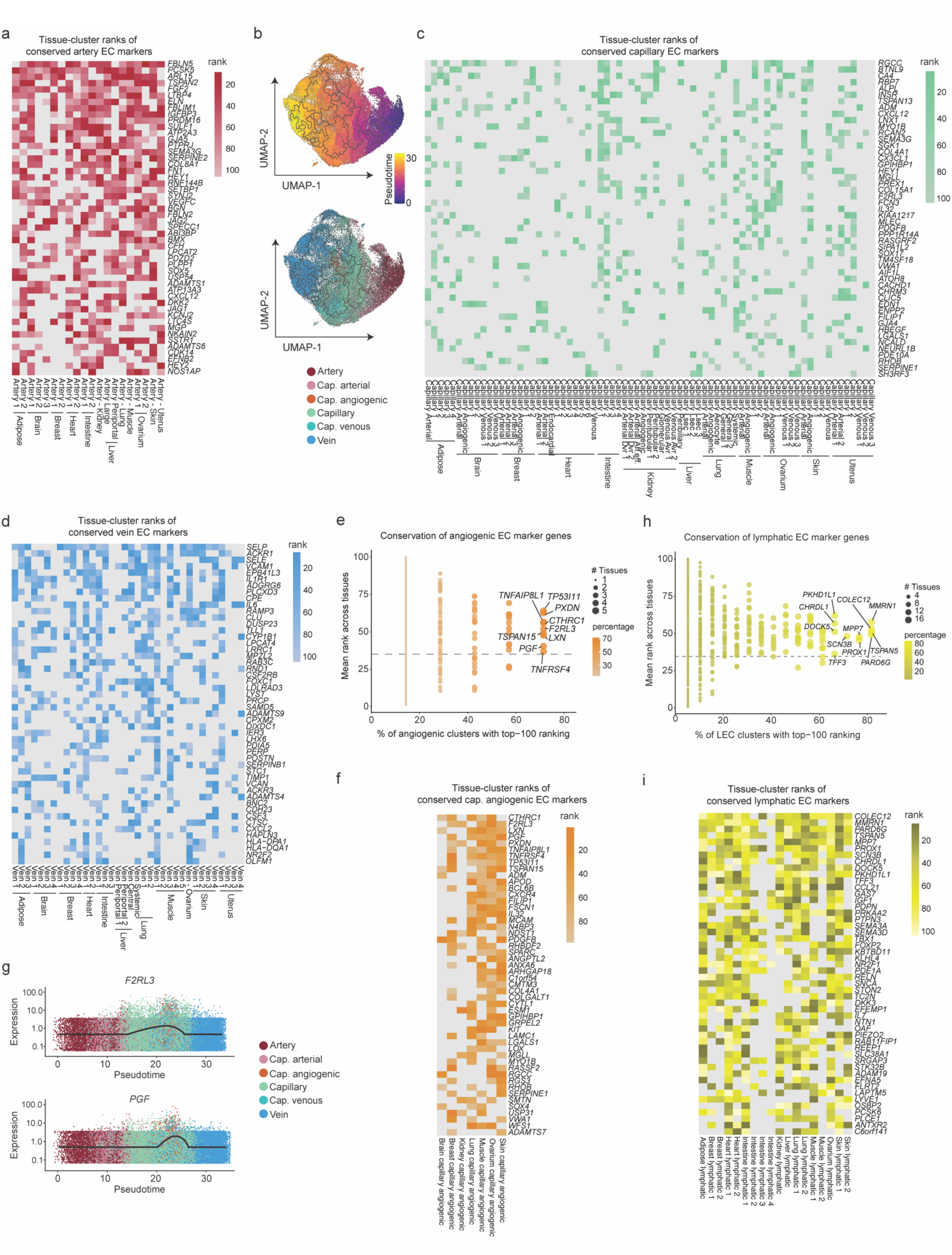
Top ranking conserved global vascular bed EC subtype marker genes. **a.** Heatmap visualization of cluster ranks of every indicated marker gene (y-axis) in artery EC subclusters (x-axis). The top 50 most highly ranked and conserved markers are shown. Grey indicates NA (i.e., gene not detected in tissue cluster). **b.** UMAP visualization of predicted pseudotime (upper panel) and global EC subclusters (lower panel). A subset/reduced version of the atlas was used for the analysis shown (see Methods). 1 indicates the start of the trajectory. **c,d.** Heatmap visualization of cluster ranks of every indicated marker gene (y-axis) in every indicated tissue-specific EC subcluster (x-axis). The top 50 most highly ranked and conserved markers are shown capillaries (c) and veins (d). Grey indicates NA (i.e., gene not detected in tissue cluster). **e.** Dot plot of conserved and tissue-specific markers in angiogenic capillary ECs. The color intensity and size of each dot represent the fraction (%) and exact number of tissues in which each respective marker gene is found in the top 100 most highly enriched genes, respectively. The top ten most conserved markers are indicated in the plot. **f.** Heatmap visualization of cluster ranks of every indicated angiogenic EC marker gene (y-axis) in every indicated tissue-specific EC subcluster (x-axis). The top 50 most highly ranked and conserved markers are shown. Grey indicates NA (i.e., gene not detected in tissue cluster). **g.** Expression of representative conserved angiogenic EC marker genes along the artery-capillary-vein axis in pseudotime. Color-coding reflects vascular bed subtype. **h.** Dot plot of conserved and tissue-specific markers in lymphatic ECs. The color intensity and size of each dot represent the fraction (%) and exact number of tissues in which each respective marker gene is found in the top 100 most highly enriched genes, respectively. The top ten most conserved markers are indicated in the plot. **i.** Heatmap visualization of cluster ranks of every indicated lymphatic EC marker gene (y-axis) in every indicated tissue-specific EC subcluster (x-axis). The top 50 most highly ranked and conserved markers are shown. Grey indicates NA (i.e., gene not detected in tissue cluster).

**Figure S3.**
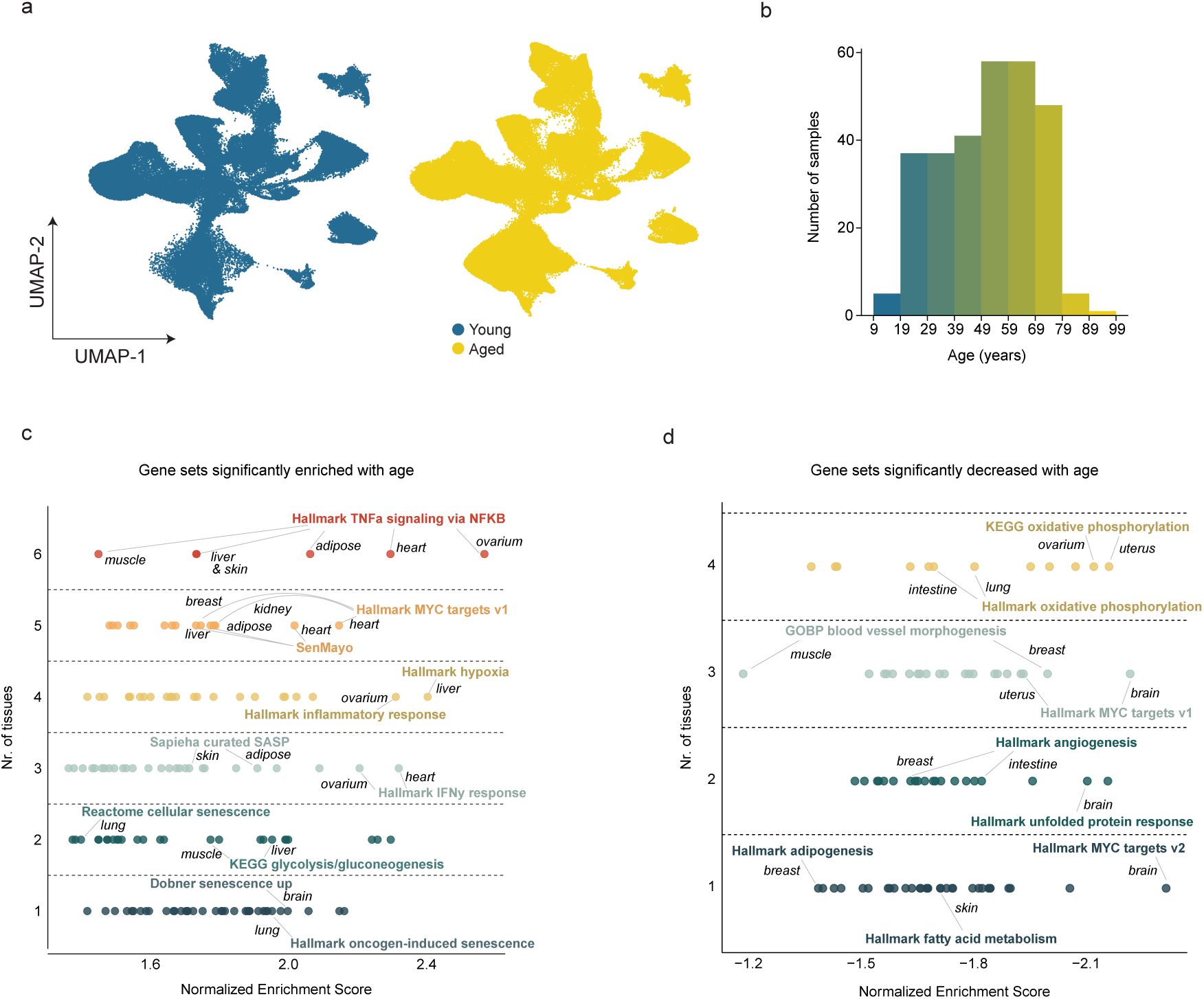
Global aging-associated EC transcriptome heterogeneity. **a.** UMAP representation of all ECs, split by age bracket. Color-coding reflects young (blue) and aged (yellow) subgroups. **b.** Bargraph of age distribution across samples in HAECA. **c,d.** Gene sets most enriched (c) and depleted (d) in aged vs. young ECs. Y-axis indicates the number of tissues in which annotated gene sets were found altered (adjusted p value < 0.25). X-axis indicates the Normalized Enrichment Score (NES).

**Figure S4.**
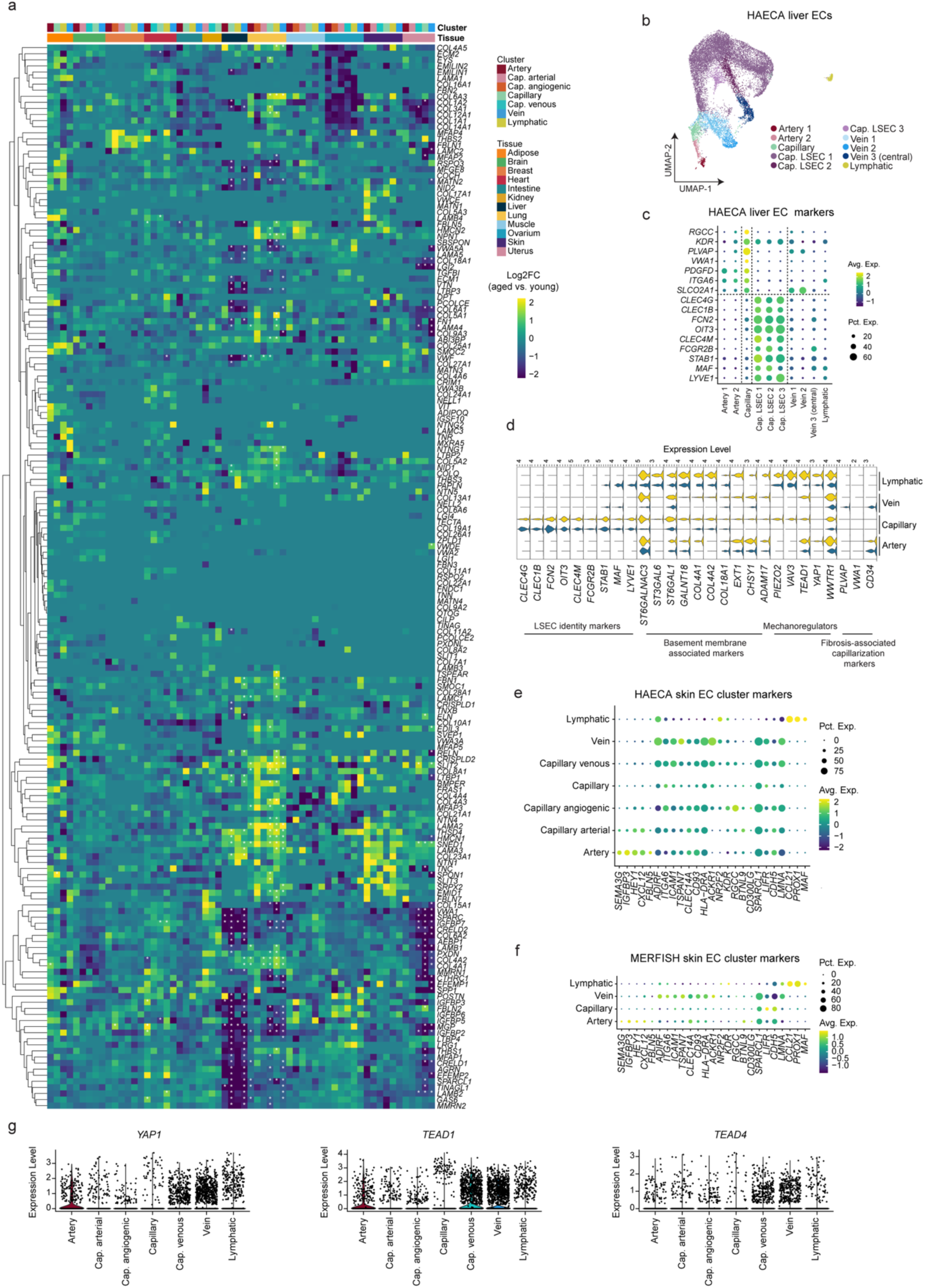
ECM-rewiring and altered mechanosignaling in the aging endothelium. **a.** Log2 fold change (Log2FC) heatmap of marker genes related to ECM and glycoproteins, per tissue and vascular bed subtype. Color scale: yellow, increased expression in aged versus young ECs; blue, decreased expression in aged versus young ECs. Asterisks indicate adjusted p-value of 0.1 or lower. **b.** UMAP representation of human liver ECs. Color-coding reflects fine EC subclusters. **c.** Dot plot heatmap of capillary- and LSEC-specific marker genes in the liver dataset. The color intensity of each dot represents the average level of marker expression, the dot size reflects the percentage of liver ECs expressing the marker within the subcluster. **d.** Violin plots of the expression of marker genes representative of LSEC identity, basement membrane regulators, mechanosensors, and fibrosis-associated capillarization, in the indicated conditions and clusters in the liver dataset. **e-f.** Dot plot heatmap of global EC subcluster markers in the HEACA skin (e) and MERFISH skin (f) EC datasets. The color intensity of each dot represents the average level of marker expression, the dot size reflects the percentage of skin ECs expressing the marker within the subcluster. **g.** Violin plots of the expression of mechanoregulatory TFs *YAP1* (left), *TEAD1* (middle) and *TEAD4* (right) in the HAECA skin dataset.

**Figure S5.**
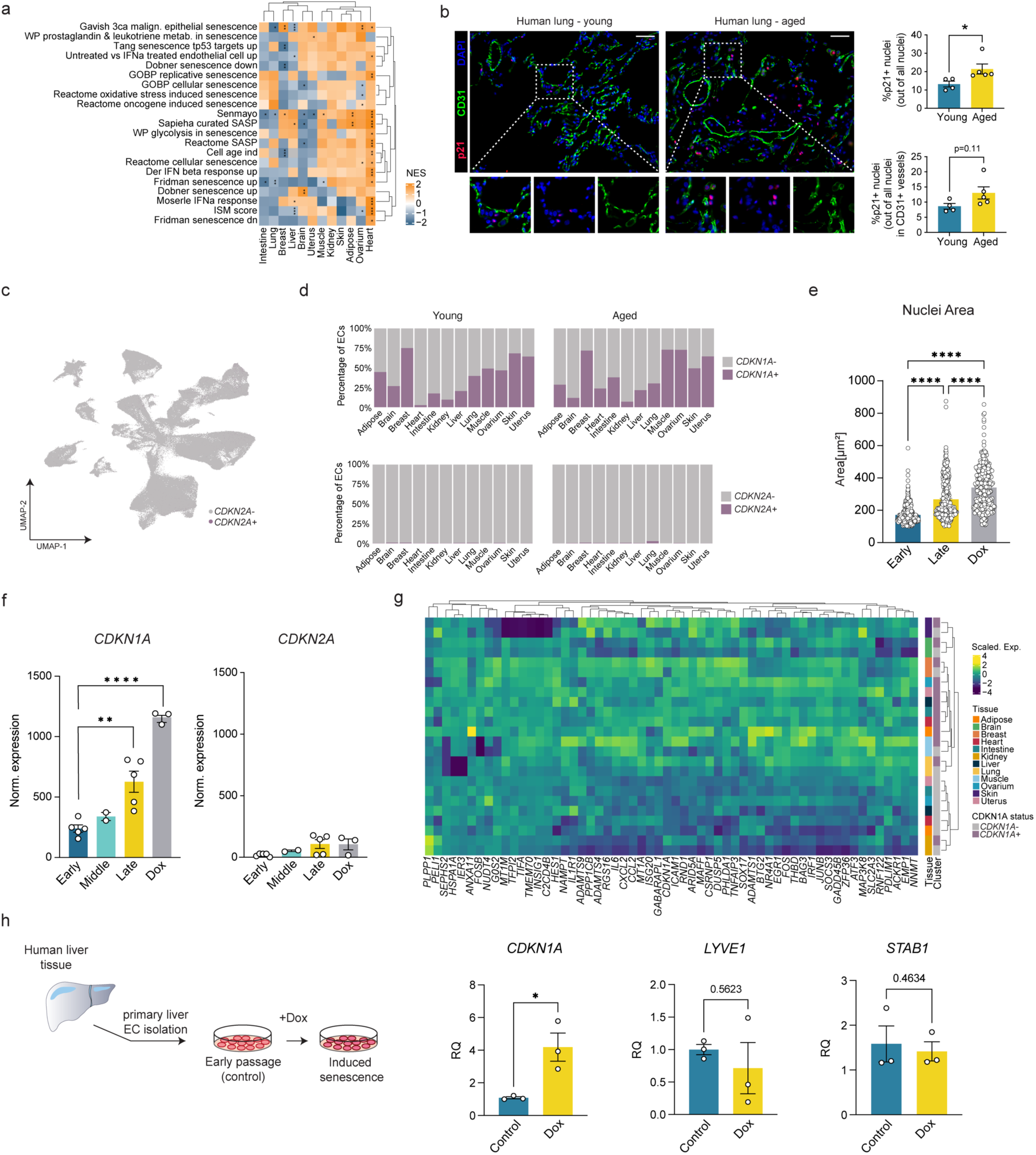
A transcriptome signature of endothelial cell senescence. **a.** Heatmap visualization of GSEA for a curated list of senescence-associated gene sets in aged ECs across tissues (compared to their young counterparts). Color scale: orange, high NES; blue, low NES. *adj.p < 0.05, **adj.p <0.01, ***adj.p <0.001. **b.** Left: Representative immunofluorescent images of human lung sections from young and aged subjects, immunostained for p21 (red), CD31 (green), and DAPI (blue). Smaller images are magnifications of the respective boxed areas. Scale bar: 50 μm. Right: Quantification of the percentage of p21-positive nuclei (top) or p21-positive nuclei in the CD31+ vessel areas (bottom) in young and aged subgroups. Mean ±SEM, unpaired t-test, two-tailed, *p < 0.05, n = 4 and 5 for young and aged groups, respectively. **c.** UMAP representation of all HAECA ECs. Color-coding reflects CDKN2A+ (purple) and CDKN2A− (grey) cell fractions. **d.** Abundance of CDKN1A+/− and CDKN2A+/− EC fractions in each tissue, per age bracket. **e.** Quantification of nuclear size in early passage, late passage, and Doxorubicin treated HUVECs (n = 7 each). Mean ±SD, *p < 0.05, by one-way ANOVA. **f.** Normalized mRNA expression values of *CDKN1A* (left) and *CDKN2A* (right) in HUVECs across different passage numbers/conditions of induced cellular senescence, as analyzed by RNA-sequencing. Mean ±SEM, **p <0.01, ****p<0.0001, by unpaired two-tailed *t*-test. **g.** Gene-expression heatmap of top 10 CDKN1A-associated markers per tissue. Color scale: yellow, high expression; blue, low expression. **h.** Left: Schematic of primary liver EC isolation and induction of cellular senescence. Right: Relative quantifications (RQ) of the normalized mRNA expression values of *CDKN1A* (left), *LYVE1* (middle), and *STAB2* (right) in primary human liver ECs under control (early passage) and Doxorubicin induced (Dox) conditions. Mean ±SD, *p <0.05, by paired t-test, n=3 independent donors.

**Figure S6.**
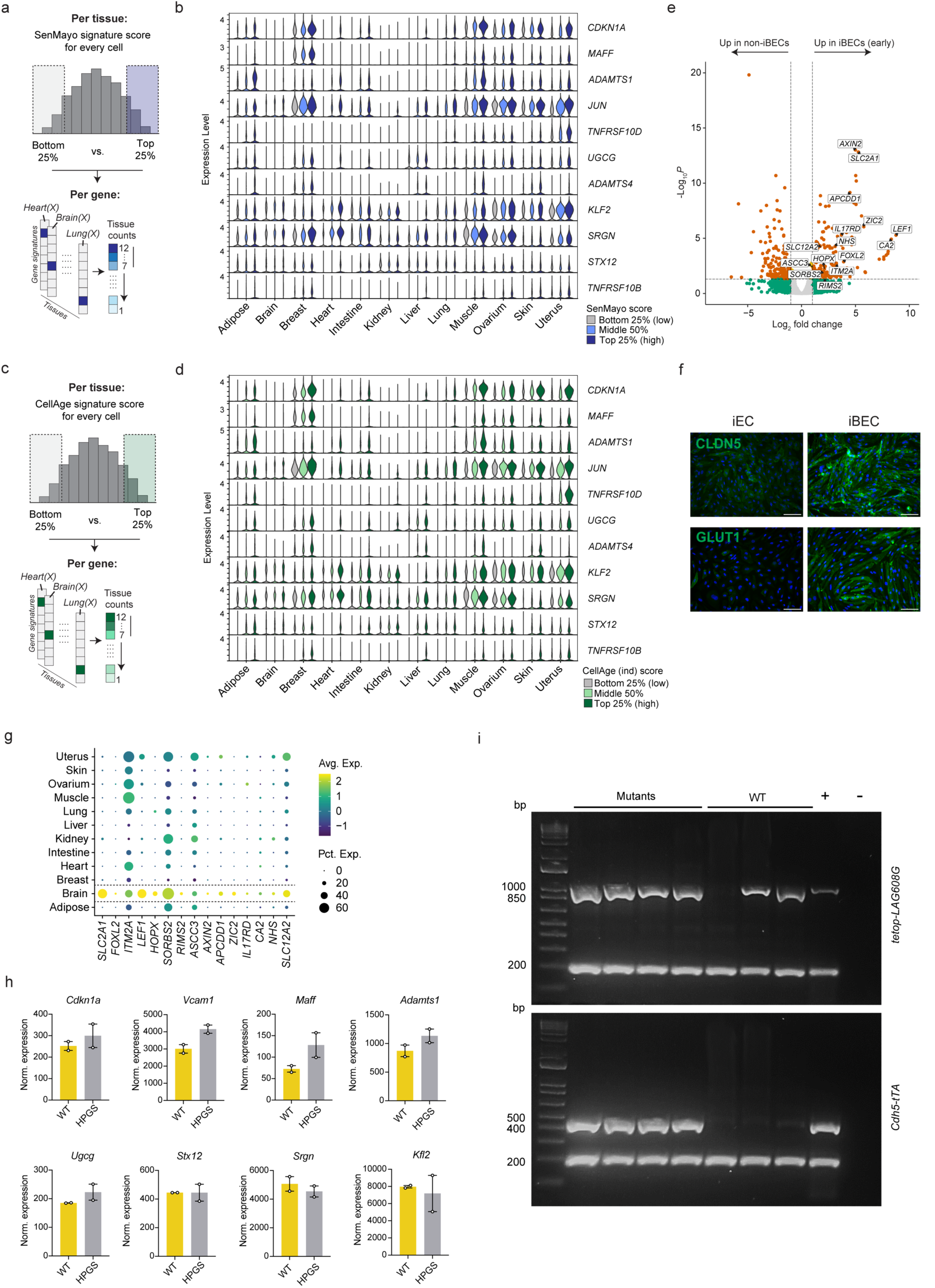
Expression of pan-tissue EC senescence markers in relation to curated senescence genesets. **a.** Schematic of SenMayo-associated analysis. **b.** Violin plots of the expression of pan-tissue senescence-associated markers in each tissue. Data is split between SenMayo^high^ (top 25%, dark blue), SenMayo^middle^ (middle 50%, light blue), and SenMayo^low^ (bottom 25%, grey) subgroups. **c.** Schematic of CellAge(ind)-associated analysis. **d.** Violin plots of the expression of pan-tissue senescence-associated markers in each tissue. Data is split between CellAge^high^ (top 25%, dark green), CellAge^middle^ (middle 50%, light green), and CellAge^low^ (bottom 25%, grey) subgroups. **e.** Volcano plot of differentially expressed genes between iBECs (right) and non-iBECs (left, iECs and HUVECs). Significant genes (adj. p-value < 0.05 and Log2 fold change > 1) are shown in orange. **f.** Representative images of CLDN5 (top) or GLUT1 (bottom) immunofluorescence staining in early passage iBECs. Nuclei are stained with DAPI (blue). Scale is 100µm. **g.** Dot plot visualization of iBEC-specific marker genes in the HAECA dataset (stratified by tissue). The color intensity of each dot represents the average level of marker expression, the dot size reflects the percentage of ECs expressing the marker within each tissue. Color scale: yellow, high expression; blue, low expression. **h.** Normalized mRNA expression values of the indicated marker genes in aortic ECs derived from WT and *Lmna^G609G/G609G^* mice (scRNA-seq dataset derived from [94]). n=2 for WT and *Lmna^G609G/G609G^* groups. **i.** Genotyping PCR for *tetop-LAG608G* (961 bp, top) and *Cdh5-tTa* (461 bp, bottom). Lower bands (∼200 bp) are internal amplification controls. Samples positive for both transgenes are classified as mutant. Samples positive for only one or neither transgene are classified as wild type. Lane 1: molecular weight DNA ladder. The last two lanes represent the positive and negative controls, respectively.

## Methods

### Human tissues

Informed consent was obtained from all research subjects. Sample collection and use were approved by the respective local ethics committees (Ethics Committee of the Medical University of Vienna (ethics approval 1969/2021), Ethics Committee Research UZ/KU Leuven (S67052), Regional Committee for Medical and Health Research Ethics, Norway (REK approval: 684971)). For more detailed donor information see **Table S10**. For lung tissues, peripheral wedge biopsies from the right middle lobe and left upper lobe lingula from donor lungs accepted for transplantation were obtained at the start of cold preservation [115]. Biopsies were fixed in 4% paraformaldehyde. Paraffin-embedded lung tissue was then sectioned at 4 µm and subjected to immunostainings as described below. For skin tissues, six-millimeter punch biopsies were obtained from skin derived from healthy individuals undergoing skin reduction surgery, embedded in optimal cutting temperature compound (OCT), snap frozen and stored at −80°C until processing. Skin biopsies were sectioned using a cryostat (Leica CM 3050 S) set at −20°C and 6 µm slice thickness. Primary human liver ECs were isolated from histologically disease-free liver tissue samples obtained during liver resections from patients undergoing surgery for non-liver primary disease.

### Animal experiments

All animal experiments were conducted in compliance with Austrian Law BGBI. I Nr. 114/2012 (TVG 2012) idF BGBI. I Nr. 76/2020 and the Guide for the Care and Use of Laboratory Animals (NIH Publication No. 85-23, revised 1996). *Prog-Tg* mouse breeding and experiments were approved (No: 2021-0.873.416) by the Ethics Committee for Laboratory Animal Experiments at the Medical University of Vienna and the Austrian Ministry of Science, Research and Economy (BMWFW-66.009/0321-WF/V/3b/2016 and BMWFW-66.009/0156-WF/V/3b/2017).

### Collection of publicly available scRNA-seq and snRNA-seq datasets

Through comprehensive literature/repository mining, we collected ECs from 59 publicly available studies. We included datasets meeting the following criteria: (1) ECs were captured; (2) donor age was specified for each sample; (3) donor/sample-specific cells were identifiable in the data; (4) raw (unnormalized) sequencing counts were provided; and (5) cells were isolated from healthy or non-pathological tissues. We excluded studies in which ECs were not clearly identifiable or captured at insufficient numbers (<350 cells per study) after pre-processing and quality control (see below). Furthermore, studies using donor pools were excluded from the analysis. We extensively harmonized available donor/sample metadata information, including age, sex, and sample collection across all studies. The final list of studies and datasets (n=34) incorporated in this study is provided in **Table S1**.

### Data pre-processing

Raw or preprocessed (but unnormalized) data were harmonized and processed in R (version 4.3.3), using the Seurat package for all following single-cell level processing steps (version 5.1) [116]. We standardized data formats by collecting mapped counts in formats such as CellRanger outputs (.features, .barcodes, .matrix files), .RDS, .H5AD, or .csv matrices. For datasets lacking GRCh38 alignment (HAECA_47), we remapped raw data using CellRanger (version 8.0.1, GRCh38.p14 (release 44)). Quality control steps for individual datasets included removing cells/nuclei with fewer than 200 unique features, outliers with high feature counts (potential doublets), and cells with >10-15%, or nuclei with >5-10% mitochondrial gene expression. For the latter two, optimal thresholds were determined per dataset via visual inspection of feature counts and calculated mitochondrial fractions. The remaining data were normalized (*NormalizeData*), and the top 2000 highly variable genes were identified (*FindVariableFeatures*). Data scaling (*ScaleData*) and dimensionality reduction (PCA; *RunPCA*) followed, with the top 25-35 PCs (determined per study) used to construct a shared nearest-neighbor (SNN) graph (*FindNeighbors*). Louvain Clustering was performed using *FindClusters* (resolution optimized per study, ranging from 1-2) and visualized using uniform manifold approximation and projection (UMAP; *RunUMAP*). EC clusters were manually identified based on canonical marker gene expression (e.g., *PECAM1, CDH5, FLT1, VWF, KDR, RGCC* for blood vessel ECs; *MMRN1, PROX1, PDPN, LYVE1* for lymphatic ECs) and lack of expression of other cellular lineage markers (e.g., immune cells *PTPRC*, stromal cells *PDGFRB*, epithelial cells *EPCAM*). All annotated EC clusters were extracted for downstream analysis.

### EC integration and annotation of vascular bed subtypes per tissue

Selected ECs from different studies were combined in a tissue-specific manner, followed by individual rounds of subclustering and annotation. As donor-specific batch effects were apparent in every combined tissue EC dataset, we explored various batch correction/data alignment methods (canonical correlation analysis (CCA), Harmony-based integration and anchor-based canonical correlation analysis (rPCA)). As rPCA resulted in the best alignment of datasets across donors and studies, we choose this method for all datasets. Briefly, the data was divided by donor (donors with <100 cells were removed from the dataset at this stage), followed by individual normalization, identification of the top 2000 highly variable genes, identification of data anchors (*FindIntegrationAnchors*) and integration of the datasets (*IntegrateData*), using the top 30 principal components. The resulting integrated data was then scaled, summarized by PCA, subclustered and annotated. Marker genes for each cluster were calculated (*FindAllMarkers,* Wilcoxon rank sum test using the “RNA” assay) and screened for coherent enrichment of canonical marker genes, to annotate artery (*GJA4, GJA5, FBLN2, FBLN5, HEY1, DKK2, CXCL12, GLUL*), capillary (*RGCC, KDR, FLT1, CA4, BTNL9*), angiogenic (*ANGPT2, ESM1, PGF, APLN, LXN, INSR*), vein (*VCAM1, SELE, SELP, ACKR1, NR2F2, HDAC9*) and lymphatic (*PROX1, PDPN, MMRN1, CCL21*) EC subtypes (**Table S2**). Capillary-arterial and capillary-venous clusters were defined as intermediates, expressing both capillary and artery- or vein markers, respectively. Specialized tissue-specific EC subtypes were furthermore annotated based on reported gene sigantures (e.g., *CLEC4G, CLEC4M, STAB2* (liver sinusoidal ECs; LSECs [39, 117]), *EMCN, PLAT* (glomerular ECs) [118]).

At this stage, in every tissue contaminating clusters were identified, either expressing non-EC marker genes (e.g., tissue-specific stromal, immune, and/or epithelial cells), low unique feature counts (likely representing remaining low-quality cells or empty droplets), and/or relatively high unique feature counts in addition to expression of both non-EC and EC markers (likely representing doublets). For the latter, we independently verified in multiple tissues that clusters assigned by us as potential doublets also harbored high predicted doublet scores using scDoubletFinder (version 1.16.0). All contaminating, low-quality, and doublet clusters were removed from further analysis. Our final dataset comprises 375,829 cells from 12 tissues (Adipose, Brain, Breast, Heart, Intestine, Kidney, Liver, Lung, Muscle, Ovarium, Skin and Uterus), 34 studies and 290 donors (**Table S3**).

### Gene filtering

For marker calculation and further analysis, ambient genes, ribosomal and mitochondrial genes were removed from the datasets by using the following pattern for exclusion: (RPS|RPL|MT-)|AS1|ENSG0000|LINC|MIR|RF000. In addition, the data was filtered for protein-coding genes, using the gene biotype “protein_coding” from the biomaRt database. Only genes detected in each study (HAECA_ID) within each tissue were retained for further analysis. Ambient RNA contamination, which can be derived from various sources (e.g., ruptured or dying cells, cell-free RNA in the cell suspension, or cell types more abundant in the cell suspension), can lead to biological misinterpretation of single-cell datasets when not properly accounted for [119, 120]. In line with this, for certain tissues in our atlas we could observe gene expression patterns indicative of likely contamination by ambient RNA derived from tissue-specific epithelial cells (e.g., *SFTPC* and *ALB* expression in a large fraction of lung and liver ECs, respectively). However, given that in our data preprocessing workflow we largely worked with mapped, raw counts data, most community-developed and well-accepted methods for detected and removal of/correction for ambient RNA contamination (e.g., CellBender, SoupX) could not be effectively used. Therefore, we addressed potential contamination from ambient mRNAs as follows: for every tissue, we analyzed the detected genes in droplets at the tail-end of the total UMI count distribution across selected HAECA_IDs (on average 50 UMIs per barcode). These likely represent empty droplets/beads instead of cells/nuclei, and hence their gene expression content is likely to represent ambient RNA contamination. We supplemented the resulting list of genes with additional genes/gene families not known to be expressed in ECs, and previously reported to likely represent contaminating ambient RNA. In total, this approach resulted in a set of 115 genes (**Table S12**) we considered a putative ‘ambient mRNA’ contamination, which we filtered out from all marker gene analyses and differential gene expression analyses in our manuscript.

### Merging and integration of pan-tissue EC atlas

Seurat objects from all integrated tissues were merged, normalized and scaled before running PCA and clustering. *FindAllMarkers()* was used to identify tissue- and subtype-specific markers. For visualization purposes only, the global atlas was integrated as described above (rPCA), once by Sampling (single-cell and single-nuclei) to visualize tissue-specific variation, and once by Tissue, to visualize vascular bed subtype specific variation of ECs.

### Congruent marker gene and trajectory analysis

For each fine EC subcluster across all tissues, we identified the top 100 positively enriched marker genes using the *FindAllMarkers()* function in Seurat (parameters: only.pos = TRUE, min.pct = 0.15, logfc.threshold = 0.25). Only genes with an adjusted p-value < 0.05 were retained, and the top markers were ranked by descending average log2 fold change (avg_log2FC). For each subcluster and tissue, we computed the frequency with which each gene appeared among these top 100 markers. To assess gene-level conservation across related endothelial subtypes, we grouped clusters based on keywords in their annotations (“artery”, “capillary”, “vein”, “lymphatic”, and “angiogenic”) and calculated gene recurrence and ranking consistency across clusters within each group. For the “capillary” analysis, arterial, venous and angiogenic capillaries were also included. For “artery” and “vein” analysis, arterial and venous capillaries were not included.

### Pseudotime analysis

Trajectory/pseudotime analysis was performed using Monocle 3 (v1.3.7). We used the global atlas dataset integrated by Tissue, and only focused on blood vascular ECs (lymphatic ECs were excluded) and data derived from single-cells to avoid cell vs. nuclei-related batch effects in the analysis. The dataset was subsampled to 5000 cells per tissue.

### Analysis of age-associated gene expression changes

For analysis of gene expression change over age, donors were binned into the age brackets “young” and “aged” depending on the median age of the donors in each tissue, ensuring balanced age brackets for downstream analyses. If the median age was > 60 years, this was set as the cutoff for age bracket binning (**Table S3**).

To allow for robust testing of differential gene expression across the different datasets, we used DESeq2 on pseudo-bulk samples as described [121]. Briefly, we aggregated gene counts of 1.) all ECs from the same sample/donor (in case of pan-EC aging signatures), or 2.) ECs from a vascular bed subtype (i.e., artery ECs, vein ECs) from the same sample/donor (in case of vascular bed EC aging signatures), using *AggregateExpression()* in Seurat on the RNA assay. Pseudo-bulk samples consisting of fewer than 10 cells were discarded from downstream analysis. Moreover, comparative analyses were only performed in case of ≥ 4 samples/donors per age bracket. We compared gene expression signatures between young and aged brackets using the Wald test (DESeq2, alpha = 0.1), including the sampling method (i.e., cells or nuclei) and the study (HAECA_ID) as a covariate. To identify congruent age-associated marker genes across tissues, we applied robust rank aggregation (RRA) using the Wald test results (DESeq2, alpha = 0.1 (Log2 fold change >0 and <0 for age-increased and −decreased genes, respectively) and the RobustRankAggreg package (v1.2.1). Benjamini-Hochberg adjusted p-values were calculated using the qvalue R package (v.2.36.0).

### Visualization of results

Visualizations were performed with ggplot2 (version 3.5.1, Wickham H (2016). *ggplot2: Elegant Graphics for Data Analysis*. Springer-Verlag New York., https://ggplot2.tidyverse.org), ggExtra (version 1.10.1, https://github.com/daattali/ggExtra), and tidyplots (version 2.0, Engler, iMeta, 2025, doi/10.1002/imt2.70018) package. *DotPlot()* and *VlnPlot()* were used for dot- and violin plot visualization of marker genes, respectively, and *DimPlot()* and *FeaturePlot()* were used for UMAP visualization of single-cell data.

### Analysis of CDKN1A senescence-associated genes

For every tissue, *CDKN1A^+^* and *CDKN1A^−^*ECs were selected (expression > 0) and annotated as senescent and non-senescent, respectively. For every HAECA_ID in a tissue, we calculated enriched marker genes (*FindMarkers()*, comparing *CDKN1A^+^* to *CDKN1A^−^* ECs, min.pct = 0.1, only.pos = TRUE). Genes enriched in at least 65% (i.e., appr. 2/3) of the HAECA_IDs in every tissue were next intersected with a list of genes significantly enriched in RNA-seq data derived from replicative senescent HUVECs (comparing late passage (P, P20-22) to early passage (P4-5)), to identify tissue-specific senescence-associated genes. From the HUVEC RNA-seq dataset, we only considered differentially expressed genes with baseMean >30, log2FoldChange > 0.35, and padj < 0.05. Next, resulting lists of tissue-specific senescence-associated genes were intersected, and genes detected in at least 7 tissues were considered robust EC senescence-associated candidate genes.

Similar analyses were performed using the SenMayo [74] and CellAge (inducing) [93] gene signatures. When present (in case of CellAge), *CDKN1A* was removed from these signatures. A senescence score per cell (separately for the SenMayo and CellAge signatures) was calculated using the *AddModuleScore()* function in Seurat, followed by categorization of ECs in High, Middle, or Low expression groups based on the upper and lower quartiles of the senescence score distribution (performed per HAECA_ID, to account for study/dataset-specific variations in expression). Within each HAECA_ID, differential expression analysis was performed using Seurat’s *FindMarkers()* function to identify genes significantly upregulated in SenMayo/CellAge^high^ versus SenMayo/CellAge^low^ cells (adjusted p < 0.05, min.pct = 0.1). For each tissue, genes consistently enriched in ≥65% of HAECA_IDs were retained as robust markers.

### Transcription Factor inference analysis

The DecoupleR package (version 2.8) [122] was used for prediction of transcription factor activities in young and aged subgroups, by using the Univariate Linear Model (*run_ulm()*) method applied to the pseudobulk Wald test results. Benjamini-Hochberg adjusted p-values were calculated using the qvalue R package.

### Gene set and GO term enrichment analyses

Gene sets were acquired from the Molecular Signature Database (MsigDB), from literature review, and from *in vitro* in-house data, and categorized into hallmark, metabolic, senescence and mechanoregulatory biological processes (**Table S7**). GSEA and GO enrichment analysis was performed as implemented in the clusterProfiler package (version 4.10.1) [123]. GO enrichment results were summarized using REVIGO [124]. Heatmap visualization of normalized enrichment scores (NES) and adj. p-values was performed using the ComplexHeatmap R package.

### Analysis of aortic ECs from *Lmna^G609G/G609G^* and WT mice

For analysis of ECs derived from the scRNA-seq dataset from [94], raw data were downloaded from ArrayExpress (EMTAB-13678), and processed and annotated as described above. ECs were subsetted based on expression of canonical markers (e.g., *Pecam1, Cdh5, Vwf, Kdr, Rgcc, Mmrn1, Prox1, Pdpn*). For analysis of *Lmna^G609G/G609G^* versus WT ECs (n=2434 ECs in total, of which 1109 WT and 1325 mutant), we created pseudobulk samples (aggregation by orig.ident and genotype) and calculated DEGs between the two conditions using the Wald test (DESeq2).

### MERFISH spatial and single-cell analysis

#### MERFISH panel design

A custom MERFISH gene panel targeting 453 genes (CP1623) (**Table S12**), plus 15 blanks to control for unspecific binding of probes, was designed using the MERFISH gene panel design tool (https://portal.vizgen.com) according to the instructions listed in the Gene Panel Design Portal manual (Vizgen, 91600101). Selection of target genes was manually curated and based on canonical as well as tissue-specific marker genes for most cell types found in human skin including T cells, macrophages, dendritic cells, Langerhans cells, endothelial cells, fibroblasts and keratinocytes. We furthermore included target genes specific for subclasses of major cell types to determine their tissue-residency, activation/exhaustion status, proliferation or response to mechanical stimuli. We evaluated the created gene panel for suitability of MERFISH imaging in terms of sufficiently long reads for probe binding and transferred six high-abundance genes to the sequential panel. To prevent optical crowding for MERFISH imaging, we confirmed that the entire gene panel meets the abundance threshold of ≤200 fragments per kilobase of transcript per million mapped reads (FPKM) for individual genes, ≥30 target regions per transcripts, <12000 FPKM for the entire panel.

#### MERFISH tissue collection, cryosectioning and permeabilization

Six-millimeter punch biopsies were obtained from skin derived from healthy individuals undergoing skin reduction surgery, embedded in optimal cutting temperature compound, snap frozen and stored at −80°C until processing. Skin biopsies were sectioned using a cryostat (Leica CM 3050 S) set at −20°C and 10 µm slice thickness and placed on a MERSCOPE slide (Vizgen, 10500001). Post sectioning, the tissue was dried for 30 minutes at −20°C and then fixed for 30 minutes at 47°C using 4% Paraformaldehyde (PFA) pre-warmed to 47°C. This was followed by three washing steps using PBS and a 1-hour drying period at room temperature, after which the sample was immersed in 70% ethanol and incubated over night at 4°C for permeabilization.

#### MERFISH sample preparation

Following overnight permeabilization, samples were processed for cell boundary staining according to the MERSCOPE User Guide (Vizgen, 91600002) using the MERSCOPE Cell Boundary Stain Kit (10400118). Briefly, blocking of unspecific binding was carried out using Blocking Buffer C, followed by a 1 hour incubation with the Cell Boundary Stain Mix (Vizgen, 20300010) and followed by a 1 hour incubation with Cell Boundary Secondary Stain Mix (Vizgen, 2030011). Following incubation, the tissue was fixed using 4% PFA and further processed for encoding probe hybridization. All steps were carried out in the presence of RNAse inhibitor (NEB. 0314S) with in-between washing steps using PBS.

#### MERFISH encoding probe hybridization

Samples were first rinsed with Sample Prep Wash Buffer (Vizgen, 20300001), then incubated at 37 °C in Formamide Wash Buffer (Vizgen, 20300002), followed by incubation with 50 µL of the custom MERSCOPE Gene Panel Mix (Vizgen, catalog no. 20300008; see **Table S12**). The tissue was covered with Parafilm to minimize evaporation and incubated in a humidified incubator set to 37 °C for 72 hours. Following this incubation, the sample was washed twice with 5 mL of Formamide Wash Buffer (Vizgen,20300002) at 47 °C for 30 minutes each, followed by a final wash with 5 mL of Sample Prep Wash Buffer for 2 minutes.

#### MERFISH gel embedding and tissue clearing

Gel embedding and tissue clearing was conducted according to the MERSCOPE User Guide (Vizgen, 91600002) with minor adaptations. Gel coverslips (PN 30200004) were prepared by first cleaning and spraying the coverslip with RnaseZap and 70% ethanol and then applying 100 µL of Gel Slick Solution (VWR, 12001-812) on top of the coverslip to be evaporated for 10 minutes at room temperature. Coverslips were used immediately after preparation. Embedding solution consisting of 5 mL Gel Embedding Premix (Vizgen, 20300004) mixed with 25 µL 10% ammonium persulfate (Millipore-Sigma, 09913-100G) and 2.5 µL N,N,Nʹ,Nʹ-tetramethylethylenediamine (Sigma, catalog no. T7024-25ML) was prepared and immediately used. Samples were embedded by applying 100 µL of the embedding solution on top of the tissue followed by overlay with the gel slick-covered coverslip and incubation for 1.5 hours to allow for polymerization of the gel. Polymerization was monitored by observing the the remaining gel embedding solution in a 50 mL test tube and considered complete once most of the gel embedding solution has solidified in the tube. Following polymerization, coverslips were carefully removed to avoid lifting of the gel and tissue digestion using Digestion Premix (Vizgen, 20300005) supplemented with RNAse inhibitor (NEB, M0314L) and an incubation of 6 hours at 37°C was conducted. Following tissue digestion, tissue clearing was conducted by first incubating the sample for 4 hours at 47°C in the clearing solution consisting of 200 µL Clearing Premix (Vizgen, 20300114) supplemented with 50 µL Proteinase K (NEB, 8107S). The clearing solution was exchanged after the incubation for a fresh solution consisting of 500 µL Clearing Premix (Vizgen, 20300114) supplemented with 50 µL Proteinase K (NEB, 8107S) and incubated overnight at 47°C after which the sample was incubated at 37°C for two days. Post-clearing, autofluorescence quenching of samples was carried out by an 8-hour incubation in the MERSCOPE Photobleacher (Vizgen, 10100003) followed by DAPI and polyT staining according to the MERSCOPE Instrument User Guide (Vizgen, 91600001). Briefly, clearing solution was aspirated and samples washed twice using Sample Prep Wash Buffer (Vizgen, 2030001), followed by a 15-minute incubation in the DAPI and PolyT Staining Reagent (Vizgen, 20300021) on a rocker. This was followed by a 10-minute incubation in Formamide Wash Buffer (Vizgen,20300002) and a washing step using Sample Prep Wash Buffer (Vizgen, 2030001).

#### MERFISH imaging

While conducting DAPI and polyT staining of the sample, the 500-gene MERSCOPE imaging cartridge (Vizgen, 20300019) was thawed in a 37°C waterbath for 1 hour. Following this, the cartridge was carefully cleaned, ten-times inverted and pierced to allow for addition of the imaging activation mix consisting of 250 µL Imaging Buffer Activator (Vizgen, 20300022) and 100 µL RNAse inhibitor (NEB, M0314L) followed by careful overlay of 15 ml Mineral oil (Millipore-Sigma, M5904-6X500ML). The flow cell was then assembled according to the MERSCOPE Instrument User Guide (Vizgen, 91600001) and MERSCOPE acquisition using our custom 500-plex gene panel CP1623 started by first conducting a low-resolution DAPI overview to specify the regions of interest which were then aquired with high-resolution spatial transcriptomic MERFISH imaging.

### Hematoxylin and eosin (H&E) imaging

For H&E staining a consecutive 6µm tissue section of each skin biopsy was obtained using a cryostat (Leica CM 3050 S). Staining and mounting of the skin sections was performed automatically using the DAKO CoverStainer. Images were acquired using a Z1 Axio Observer microscope equipped with a LD Plan-Neofluar 20x/0.4 objective (Zeiss).

### Spatial analysis of MERFISH dataset

Stitched image files were segmented by the Vizgen Post-processing Tool (vpt) version 1.2.2 with the CellPose cyto2 model. Resulting cell segmentation masks were visually inspected to confirm correspondence with nuclear and cytoplasmic stain. Count matrices of transcripts per cell as created by vpt were used for downstream analysis using the scverse tool suite. Cells containing at least 10 transcripts were retained in the dataset. Spatial plots were created using Scanpy (*pl.spatial*).

### Single-cell analysis of MERFISH dataset

Single-cell data from MERFISH experiment was processed in R (version 4.3.3), using the Seurat package (version 5.1) [116]. The data was normalized (*NormalizeData*), and the top 2000 highly variable genes were identified (*FindVariableFeatures*). Data scaling (*ScaleData*) and dimensionality reduction (PCA; *RunPCA*) followed to construct a shared nearest-neighbor (SNN) graph (*FindNeighbors*). Louvain Clustering was performed using *FindClusters* (resolution = 1) and visualized using uniform manifold approximation and projection (UMAP; *RunUMAP*). EC clusters were identified based on canonical marker gene expression (e.g., *PECAM1, CDH5, FLT1, VWF, KDR, RGCC* for blood vessel ECs; *MMRN1, PROX1, PDPN, LYVE1* for lymphatic ECs) and lack of expression of other cellular lineage markers (e.g., immune cells *PTPRC*, stromal cells *PDGFRB*, epithelial cells *EPCAM*). EC clusters were extracted for downstream analysis of subclustering and annotation. Normalization, scaling, summarized by PCA and dimensionality reduction was repeated with the EC subset, as described above. Marker genes for each cluster were calculated screened for coherent enrichment of canonical marker genes, to annotate artery (*GJA4, GJA5, FBLN2, FBLN5, HEY1, DKK2, CXCL12, GLUL*), capillary (*RGCC, KDR, FLT1, CA4, BTNL9*), angiogenic (*ANGPT2, ESM1, PGF, APLN, LXN, INSR*), vein (*VCAM1, SELE, SELP, ACKR1, NR2F2, HDAC9*) and lymphatic (*PROX1, PDPN, MMRN1, CCL21*) EC subtypes. Genes of interest were highlighted using the *FeaturePlot* function.

### EC-Progeria mouse model and EC isolation

#### Generation of bitransgenic mouse lines

Bi-transgenic *Prog-Tg* we generated *de novo* by crossing tet-operon-driven transgenic mice (C57BL/6J background) carrying the HGPS mutant *lamin A* minigene (1824C>T; G608G; *tetop-LAG608G*) with transgenic mice expressing a tetracycline-responsive transcriptional activator under the endothelial cell-specific VE-Cadherin promoter (*Cdh5-tTA* mice, Jackson Laboratories MGI:4437711, FVB background) as previously described [125–127]. Animals were maintained at C57BL/6J background (N5 generation; ∼97%). Mice were maintained without doxycycline to permit continuous expression of the respective transgenes in bitransgenic animals. Genomic DNA was isolated from the distal phalanx of each mouse for genotyping. PCR amplification was performed with 1.) an initial denaturation at 94 °C for 5 min, followed by 29 cycles of 94 °C for 30 s, 59 °C for 1 min, and 72 °C for 45 s, with a final extension at 72 °C for 2 min (*lamin A*), or 2.) an initial denaturation at 94 °C for 5 min, then 29 cycles of 94 °C for 30 s, 57 °C for 1 min, and 72 °C for 45 s, followed by a final extension at 72 °C for 2 min (*Cdh5-tTA*). For primers, see **Table S12**.

### Endothelial cell isolation and RNA extraction

Primary mouse endothelial cells were isolated as previously described [125]. Briefly, hearts and livers from young animals (age 21 days) were digested with 200 U/ml collagenase I (Gibco, 17100-017), under constant rotation for 45 minutes at 37 °C The tissue digest were triturated by passage through a 19-gauge needle, and filtered through a 70 µm cell strainer to obtain single-cell suspensions. ECs were isolated by magnetic bead sorting using rat anti-ICAM2 antibodies (BD Biosciences, clone 3C4 [mIC2/4], 553326) bound to sheep anti-rat IgG–coated paramagnetic Dynabeads (Invitrogen, 11035). After extensive washing to remove non-endothelial cell populations, enriched ECs were lysed in 350 µL or 700 µL QIAzol (Qiagen, 79306) depending on tissue size. Cell extracts were stored at −80°C to preserve RNA integrity until RNA extraction.

QIAzol-samples were thawed at RT and processed immediately. Chloroform was added at 20% volume of Qiazol (70 µL or 140 µL), vortexed and incubated at room temperature (RT) for 2-3 min. Samples were centrifuged for 15 minutes at 4°C and 12.000 xg. The upper phase was harvested and transferred into a new tube. GlycoBlue carrier (1.2 µL or 2.4 µL, Invitrogen) and ice-cold isopropanol at 50% volume of Qiazol (175 µL or 350 µL) were added to the sample, vorexted and incubated at RT for 10 minutes. Samples were centrifuged for 10 minutes at 4°C and 12.000 g. Supernatant was removed and 75% Ethanol (VWR), at 100% volume of Qiazol (350 µL or 700 µL) was added to the pellet. Samples were centrifuged for 10 minutes at 4°C and 12.000 g. Supernatant was removed and pellet was dried open for 20 minutes. Pellet was resuspended in 15-30 µL of molecular grade RNAse free water, depending on the size of the tissue. RNA was quantified on a NanoDrop Photometer (Thermo Fisher) and stored at −80°C.

### Realtime PCR and analysis

Reverse transcription was performed using LunaScript RT SuperMix Kit (New England Biolabs) according to manufacturer’s protocol, using 750 ng of RNA. qRT-PCRs were performed using Luna Universal qPCR Master Mix (New England Biolabs) according to manufacturer’s protocol. cDNA was diluted 1:10, and 2 µL were used for each technical replicate, performed in triplicates. qRT-PCR was performed on a BioRad CFX96 Touch Real-Time PCR Detection System. The delta delta ct method was used to quantify gene expression relative to other samples. For primers, see **Table 12**.

### Endothelial cell isolation and culture

Human Umbilical Vein Endothelial Cells (HUVECs, donor pools, Sigma-Aldrich) were cultured at 37°C and 5% CO_2_ in Endothelial Cell Growth Medium 2 with supplements (EGM2, Sigma-Aldrich) and 1% Penicillin-Streptomycin (Sigma-Aldrich) and passaged at 80-90% confluency. Young (passage 3-5) and aged (passage 15-20, replicative senescence) HUVECs were used for immunofluorescence staining or for RNA extraction. Cells were regularly tested and found to be free of mycoplasma contamination. For Doxorubicin-induced senescence, early passage HUVECs were cultured until 90% confluence and treated with 250nM of Doxorubicin (Medchem Express) in EGM2 for 24 hours. Medium was replaced by fresh EGM2 medium, and cells were cultured for 6 more days, replacing medium every other day.

Primary human liver ECs were isolated from histologically disease-free liver tissue samples obtained during liver resections from patients undergoing surgery for non-liver primary disease (**Table S10**), with informed consent. Tissue samples were collected an minced on ice, washed with EDTA and digested with collagenase P (Sigma-Aldrich), based on a previously published protocol [128]. ECs were isolated using sequential centrifugation method, as described [129] and plated on 2,5% Geltrex (ThermoFisher Scientific) coated plates. Liver ECs were cultured at 37°C and 5% CO2 in Human endothelial SFM (ThermoFisher Scientific), supplemented with 1% FBS (ThermoFisher Scientific), 30 ng/ml VEGF (Peprotech), 20 ng/ml bFGF (Peprotech) and optionally 10 μM Y-27632 (during first 48 after isolation). For Doxorubicin-induced senescence, liver ECs were cultured until 75-80% confluence and treated with 500nM of Doxorubicin (Medchem Express) in Human endothelial SFM. After 24 hours of treatment cells were trypsinized and collected for RNA isolation using RNeasy Mini Kit (Qiagen) according to the manufacturer’s instructions.

### Induced (brain) endothelial cell (iEC and iBEC) differentiation

Induced pluripotent stem cells (iPSCs, KOLF2.1J) were cultured at 37°C and 5% CO_2_ in Gibco Essential 8 Flex Medium (E8Flex, Fisher Scientific) and passaged or differentiated at 80-90% confluency, using Gibco StemPro Accutase Cell Dissociation Reagent (Fisher Scientific) for 3-5 minutes at 37°C and pelleting at 300 xg for 5min. Cells were resuspended in E8Flex medium, supplemented with 1µM of Thiazovivin (MedChem Express), and seeded into cell-culture plates that have been pre-coated with Geltrex at 1:100 dilution. Growth medium was replaced one day after seeding.

iEC and iBEC differentiation protocols were adapted from [130, 131]. For mesoderm differentiation, growth medium was removed from iPSC cultures and cells were washed twice with PBS before addition of DMEM/F12 medium (Fisher Scientific), supplemented with 1X Gibco B-27 Supplement (Fisher Scientific), 1X Gibco N-2 Supplement (Fisher Scientific), 50 nM of 2-Mercaptoethanol (Sigma-Aldrich), 6 µM of Laduviglusib (Medchem Express) and 25 ng/mL of Bone morphogenetic protein 4 (BMP-4). Cells were cultured overnight at 37°C and 5% CO_2_, and medium was replaced with fresh DMEM/F12 medium (Fisher Scientific), supplemented with Gibco B-27 Supplement (Fisher Scientific), 1X Gibco N-2 Supplement (Fisher Scientific) and 50 nM of 2-Mercaptoethanol (Sigma-Aldrich), on the following day.

For iEC and iBEC differentiation, growth medium was removed from mesoderm cultures and cells were washed twice with PBS before addition of Gibco StemPro-34 Serum-free Medium, supplemented with 1:100 GlutaMAX supplement (Thermo Fisher Scientific), 350 µg/mL of Bovine Serum Albumin (BSA, Sigma-Aldrich), 5 µM of Forskolin (Medchem Express), 50 ng/mL of recombinant human VEGF165 protein (VEGF, Medchem Express), 50 ng/mL of Fibroblast growth factor 2 (bFGF, Medchem Express). Additionally, for iBEC differentiation, 10 µM of Retinoic Acid (RA, Medchem Express) was added to the medium. Cells were cultured for two days at 37°C and 5% CO_2_, medium was replaced by new, fully supplemented endothelial differentiation medium (respective supplements for iEC and iBEC), and cells were cultured for two additional days.

For EC selection, cells were trypsinized and harvested at 300 xg for 5 minutes. Cells were stained with 1:50 CD144-PE antibody (Miltenyi Biotec) in 0.5% BSA (Sigma-Aldrich) and 2mM EDTA (Sigma-Aldrich) in PBS. Positive, single cells were selected against unstained cells in the PE-channel on a Fluorescence-Activated Cell Sorter (FACS) (Sony LE-SH800). Sorted iECs were seeded into a cell-culture flask pre-coated with 0.1% gelatin (Sigma-Aldrich) and cultured at 37°C and 5% CO_2_ in EGM2 (Sigma-Aldrich), supplemented with 50 ng/mL of VEGF (Medchem Express). Sorted iBECs were seeded into a pre-coated cell-culture plate and cultured at 37°C and 5% CO_2_ in Gibco Human Endothelial Serum-free Medium (hESFM, Fisher Scientific), supplemented with 1X of Gibco B-27 Supplement (Fisher Scientific) and 50 ng/mL of bFGF (Medchem Express). The following day, 6 µM of Laduviglusib (Medchem Express) and 50 ng/mL of bFGF (Medchem Express) were added to the medium. After two more days of culture, growth medium on iBEC cells was replaced by fresh, fully supplemented hESFM with 6 µM of Laduviglusib (Medchem Express) and 50 ng/mL of bFGF (Medchem Express).

### SA-β-gal staining

For senescence detection, young and aged HUVECs were seeded at a density of 12.000 cells/well into a 96-well cell-culture plate, pre-coated with 80µg/ml of Collagen-I (Sigma-Aldrich) in acetic acid for 30 minutes at 37°C. Cells were washed with PBS twice, and cultured overnight. SPiDER-Βgal-FITC from the FastCellular Senescence Detection Kit (MP Biomedicals) was used as stated in manufacturer’s protocol at a 1:1000 dilution in McIlvaine buffer, containing 20 mmol/L citric acid and 40 mmol/L of sodium phosphate, adjusted to pH 6. Growth medium was aspirated from the wells and cells were fixed in 4% paraformaldehyde (Thermo Fisher Scientific) for 15 minutes at room temperature, followed by two rounds of washes in PBS. Afterwards, 50µl of diluted SPiDER-FITC staining was added to the cells, followed by a 30 minute incubation at 37°C (sealed). Wells were washed twice in PBS and imaged, or subjected to additional immunofluorescence stainings, as described below.

### Immunofluorescence

For immunofluorescence, early and late passage HUVECs were seeded at a density of 12.000 cells/well, and iECs and iBECs were seeded at a density of 20.000 cells/well, into a 96-well cell-culture plate, that has been pre-coated with 80µg/ml of Collagen-I (Sigma-Aldrich) in acetic acid for 30 minutes at 37°C and washed with PBS twice, and cultured overnight. Growth medium was aspirated from the wells and cells were fixed in 4% paraformaldehyde (Thermo Fisher Scientific) for 15 minutes at room temperature, followed by two rounds of washes in PBS. Cells were washed twice with PBS, followed by permeabilization with 100 µL of 0.1% Triton X-100 (Sigma-Aldrich) in PBS for 10 minutes at room temperature. Cells were blocked in 100 µL of 2% BSA (Sigma-Aldrich) in PBS for 60 minutes at room temperature, followed by staining with 50 µL of primary antibody (rabbit-p21 at 1:500 (Cell Signaling) for HUVECs, rabbit-GLUT1 (Invitrogen) at 1:100 or rabbit-CLDN5 (Invitrogen) at 1:300 for iEC and iBECs) overnight at 4°C. Cells were washed three times for 10 minutes in 2% BSA in PBS, followed by addition of 50 µL secondary antibodies (goat anti-rabbit AF594 (Invitrogen) for HUVECs, goat anti-rabbit AF488 (Invitrogen), all at 1:500) and DAPI at 1:500 (Invitrogen) for nuclear stain. Cells were incubated for 60 minutes at room temperature, followed by three more washes. Stained HUVECs were imaged on the EVOS M5000 imaging system (Thermo Fisher Scientific) at 20X magnification. Stained iEC and iBECs were imaged on a Leica DMI6000 B microscope (Leica microsystems) at 20X magnification.

### Immunofluorescence (human tissues)

Paraffin-embedded lung tissue sections were deparaffinized by incubation of slides at 60°C for 20 minutes, followed by washing steps in UltraClear (VWR) for 10 minutes, 100% Ethanol (VWR) for 5 minutes, followed by 2 minutes each of 96% Ethanol, 70% Ethanol and distilled water. Heat-induced epitope retrieval was performed in 10mM TRIS (VWR), 1mM EDTA (Sigma-Aldrich) and 0.05% Tween-20 (Fisher Scientific) buffer adjusted to pH 9 at 96°C for 30 minutes, followed by 5 minutes of washing in distilled water and 10 minutes in PBS. For immunostaining, deparaffinized lung tissue or thawed OCT-embedded skin tissue sections were permeabilized in 0.1% Triton X-100 (Sigma-Aldrich) in 2% BSA (Sigma-Aldrich) in PBS for 30 minutes at room temperature, followed by blocking in 3% goat serum (heat-inactivated, Rockland) in 2% BSA in PBS for 45 minutes followed by incubation in primary antibody at respective dilution (rabbit-p21 (Cell Signaling) and mouse-CD31 (Cell Signaling), both at 1:100 for lung tissues, rabbit-YAP (Invitrogen) at 1:600 and mouse-CD31 (Cell Signaling at 1:1000 for skin tissues) or with isotype control of respective species at same final concentration (Rabbit IgG Isotype Control or Mouse IgG Isotype Control, both Invitrogen)) in 3% goat serum (Rockland) and 2% BSA in PBS at 4°C overnight. Sections were washed three times for 10 minutes each in PBS before adding secondary antibodies goat anti-mouse AF647 (Invitrogen) at 1:400 and goat anti-rabbit AF546 (Invitrogen) at 1:200 for skin tissue, or goat anti-rabbit AF594 (Invitrogen) at 1:500 and goat anti-mouse AF647 (Invitrogen) at 1:500 for lung tissue, in 3% goat serum and 2% BSA for 60 minutes at room temperature, followed by three more washes with PBS as described before. For highly auto-fluorescent lung tissues, Vector TrueVIEW Autofluorescence Quenching Kit with DAPI (Szabo Scandic) was used for quenching and mounting and used per manufacturer’s instructions. For skin tissues, nuclei were stained with 1:1000 DAPI (Invitrogen) for 10 minutes at room temperature and washed once with PBS before adding Fluoromount-G Mounting Medium (Thermo Fisher Scientific 00-4958-02). Images were acquired on the TissueFAXS system (TissueGnostics, Austria) using the TissueQuest software (TissueGnostics, version 7.1) and analyzed in ImageJ and with the TissueFAXS Viewer software (TissueGnostics, version 7.1). Images from lung tissues were acquired on the EVOS M5000 Imaging System (Invitrogen, Thermo Fisher Scientific) and analyzed in ImageJ.

### RNA extraction, sequencing and analysis

For RNA preparation from cultured cells, samples were trypsinized and stored in RLT buffer. RNA was isolated using the RNeasy Mini Kit (Qiagen) according to the manufacturer’s instructions. RNA concentration and purity were accessed with NanoDrop One (Thermo Fisher Scientific, MA, United States) and submitted to the Biomedical Sequencing Facility of the CeMM Research Center for Molecular Medicine of the Austrian Academy of Sciences for further processing. NGS libraries were prepared from total RNA samples with the QuantSeq 3’ mRNA-Seq V2 Library Prep Kit with UDI (Cat. No: 191.96, Lexogen GmbH, Vienna, Austria). Resulting library concentrations were quantified with the Qubit 2.0 Fluorometric Quantitation system (Life Technologies, Carlsbad, CA, USA) and the fragment size distribution was assessed using the High Sensitivity DNA Kit (5067-4626, Agilent) on a 2100 Bioanalyzer High-Resolution Automated Electrophoresis instrument (Agilent, Santa Clara, CA, USA). Before sequencing, sample-specific NGS libraries were diluted and pooled in equimolar amounts. Expression profiling libraries were sequenced on a NovaSeq 6000 instrument (Illumina, San Diego, CA, USA) following a 50-base-pair, single-end recipe. NGS reads were mapped to the Genome Reference Consortium GRCh38 assembly via “Spliced Transcripts Alignment to a Reference” (STAR, 2.7.11b) utilising the “basic” GENCODE transcript annotation from version v46 (May 2024) as reference transcriptome. Since the hg38 assembly flavour of the UCSC Genome Browser was preferred for downstream data processing with Bioconductor packages for entirely technical reasons, GENCODE transcript annotation had to be adjusted to UCSC Genome Browser sequence region names. STAR was run with options recommended by the ENCODE project. Metadata annotation, such as the start of NGS adapter sequences (in XT tags), or unique molecular index (UMI) sequences (in RX tags) and UMI base quality scores (in QX) tags were propagated from unaligned BAM files to the aligned BAM files via Picard MergeBamAlignment and duplicated alignments were marked with Picard MarkDuplicates in a UMI-aware manner but not removed. NGS read alignments overlapping Ensembl exon features were counted with the Bioconductor GenomicAlignments (1.42.0) package via the summarizeOverlaps function in Union mode, ignoring secondary alignments, alignments not passing vendor quality filtering and for UMI-supporting protocols only, also duplicate alignments. Since the QuantSeq 3’ mRNA-seq FWD protocol leads to sequencing of the second strand, alignments were counted strand-specifically in feature (i.e., gene, transcript, and exon) orientation. Exon-level counts were aggregated to gene-level counts, and downstream analysis was performed using DESeq2.

### Image Analysis

#### Segmentation of HUVEC nuclei and MFI quantification (for p21)

All Image processing was performed in ImageJ/Fiji (Version v1.54p). For each image, DAPI stain was used for gaussian blur, thresholding and segmentation of ROI with a minimum of 100 µm^2^ area. Area and mean fluorescence intensity (MFI) of each ROI was measured for DAPI and p21 stains. MFIs were normalized to MFI of DAPI from the same region. Resulting data was analyzed in Prism (GraphPad).

### Segmentation of SA-β-gal stained cells

For each sample, a composite of the two recorded channels (DAPI and GFP) was created and scaled (0.259 µm per pixel). Voxel depth was set to 1 µm for metadata consistency. Cell segmentation was performed with the Cellpose Fiji plugin using the cyto_3 model. The model was run on the two-channel composite with the following parameters: diameter = 180, ch1 = 1 (cytoplasm/GFP) and ch2 = 2 (nuclei/DAPI); environment type venv with an existing Python environment path. The resulting Cellpose segmentation mask was converted into ROIs via Label Map to ROIs (MorphoLibJ), using C4 connectivity, vertex location = Corners, and name pattern r%03d. ROIs were displayed with outlines and labels in the ROI Manager for visual quality control. GFP channel (C1) within the Cellpose generated mask was quantified using the ROI Manager and per-object measurements (Measure).

### Analysis of p21+ cells in human tissue

DAPI, CD31, and p21 images were manually smoothed and thresholded to generate binary masks. A watershed operation was applied to the DAPI mask to segment nuclei. Binary mask images were analyzed using a custom macro. For each image set, the DAPI mask was used to identify nuclei using the “Analyze Particles” function with a size threshold of ≥50 pixels² and circularity between 0.5 and 1.0. Each detected nucleus was stored as a region of interest (ROI). p21 expression was assessed by measuring the mean intensity within each ROI on the corresponding p21 image. ROIs with a non-zero mean intensity were classified as p21-positive. Subsequently, the same p21-positive ROIs were measured on the CD31 image to determine co-localization with endothelial cells, defined by a non-zero CD31 signal. For each image set, three metrics were recorded: (1) total number of nuclei, (2) number of p21-positive nuclei, and (3) number of p21-positive nuclei co-localizing with CD31. Results were appended to a .CSV report for batch-level analysis.

### Analysis of nuclear versus cytoplasmic YAP

The skin epidermis and overexposed areas (i.e. glands) were manually excluded from analysis. The endothelial cell channel and nuclear (DAPI) images were smoothed (twice), followed by manual thresholding to generate binary masks. The DAPI mask was further refined using the watershed algorithm and additional smoothing to separate closely adjacent nuclei. An intersection (AND operation) of the EC and DAPI masks was used to define nuclear regions within ECs. Cytoplasmic regions were derived by edge detection (Find Edges) on the EC-DAPI mask, followed by Gaussian blur, auto-thresholding, and binarization. Regions of interest (ROIs) for both nuclear and cytoplasmic compartments were extracted using particle analysis with size and circularity filters, and subsequently used to quantify YAP signal intensity.

Subsequent analysis was performed in R. Nuclear and cytoplasmic YAP intensities were extracted from corresponding ROIs by matching coordinates using minimum Euclidean distance, compensating for non-sequential ROI ordering in ImageJ. Only nuclei within a defined size range and with a minimal distance from their matched cytoplasmic region (<2000 pixels) were retained. ROIs with saturated signals (mean intensity of 255) or invalid ratios (e.g., 0, NA) were excluded. The nuclear-to-cytoplasmic YAP intensity ratio was computed for each cell, and summary metrics, including the average ratio and the fraction of cells with nuclear-enriched YAP (ratio >1), were calculated for each sample.

### Quantification and statistical analysis

Statistical analyses were performed using R or GraphPad Prism (GraphPad Software, USA). For gene expression comparisons between young and aged subgroups (scRNA-seq pseudobulk samples and bulk RNA-seq), we used the Wald test (DESeq2) with multiple testing using the Benjamini and Hochberg method. To identify congruent age-associated marker genes across tissues, we applied robust rank aggregation using the Wald test results, and calculated Benjamini-Hochberg adjusted p-values (qvalue R package). For immunostaining and qPCR results, comparison of changes between two groups was performed using an unpaired, two-tailed t-test (normally determined by performing a Shapiro-Wilk test) or a Welch’s test (unpaired; two-tailed; in case data was not normally distributed). In case of >2 comparisons, one-way ANOVA was used with a post hoc Tukey test. All immunofluorescence- or histochemical analyses were repeated in a minimum of 4 donors/samples per group or condition, and representative images are displayed. Linear regression and Pearson correlation analyses were performed separately for each group to assess the relationship between candidate genes and p21 mean intensities. A minimum of 3 independent donor pools were used for all experiments involving HUVECs. iBEC results were created using 3 independent rounds of differentiation.

